# Characterisation of *cis*-regulatory elements and genome wide enhancer prediction in Heliconiini butterflies

**DOI:** 10.64898/2026.10.28.762477

**Authors:** Adam D Ritchie, Francesco Cicconardi, Stephen H Montgomery

## Abstract

The large-scale annotation and spatiotemporal characterisation of *cis*-regulatory promoters and enhancers remain limited in many non-model organisms, largely due to fragmented availability of multiomic datasets across species and developmental conditions. *Heliconius* butterflies, which have been used to investigate wing development and patterning, have provided key insights into the genetic architecture underlying morphological diversification and adaptive pigmentation. Despite this, we are limited in what we understand of how *cis*-regulatory dynamics drive phenotypic wing patterning variation. Here, by re-analysing and re-evaluating available data, we present a comprehensive new description of a *cis*-regulatory atlas for *Heliconius* wing development by integrating chromatin accessibility dynamics (ATAC-seq) with histone modification profiling (H3K27ac, H3K4me3). High-confidence peak sets enabled robust annotation of promoters and enhancers, revealing regulatory distributions consistent with canonical regulatory organisation. Promoters displayed comparatively higher conservation of transcription factor motifs, whilst enhancers exhibited greater sequence diversity, broader genomic distributions and more variable motif repertoires. Comparative analysis revealed *cis*-regulatory divergence to be primarily temporal, with substantial chromatin remodelling and differential histone enrichment across development, while regulatory dynamics were highly conserved between forewing and hindwing tissues. Extending these experimentally characterised regulatory landscapes, a convolutional neural network was trained for sequence-based enhancer prediction, achieving test AUROC and AUPRC values of 0.909 and 0.913, respectively. Predictions across 34 Heliconiini assemblies displayed significant enrichment within independently characterised accessible chromatin and conserved non-exonic elements (CNEEs), supporting their putative regulatory function. Overall, this study provides a large-scale characterisation of *cis*-regulatory dynamics and extends enhancer annotation across Heliconiini, providing a foundation for comparative investigations of regulatory conservation and diversification.

## Introduction

Developmental processes and phenotypic diversification are intrinsically linked to gene regulatory networks, in which *cis*-regulatory elements (CREs) control where, when, and to what extent genes are expressed, thereby shaping spatiotemporal patterns of gene activity and contributing to phenotypic divergence between species (Wittkopp and Kalay, 2011; Moore *et al*., 2026). Regulatory evolution has long been proposed to mediate substantial phenotypic differences despite limited divergence in protein coding sequences (King and Wilson, 1975). Subsequent comparative and functional genomic studies have provided extensive empirical support for this model in which the modular architecture of CREs enables changes in quantitative and spatiotemporal gene expression patterns while leaving protein function unchanged, thereby reducing the pleiotropic constraints associated with coding sequence evolution (Wittkopp and Kalay, 2011). A comprehensive understanding of CREs is therefore critical in elucidating how a single genome gives rise to the many genomic blueprints that determine the behaviour of a diverse range of cell types during development (Zeitlinger *et al*., 2025).

Two of the principal CREs, promoters and enhancers, act in a combinatorial fashion to influence activation of gene expression directed by their respective spatiotemporal developmental context (Galouzis and Furlong, 2022). Investigating this regulatory architecture therefore depends on the genome wide annotation and functional characterisation of promoters and enhancers, including their genomic localisation, transcription factor associations, and differential regulatory activity during development. Promoter annotation is comparatively straight forward due to their constrained positioning proximal to transcription start sites (TSSs), while enhancer annotation is substantially more challenging. Enhancers can occur within intronic or intergenic regions, often at considerable and highly variable distances from their cognate genes, and also lack a universal sequence signature (Chan *et al*., 2009; Stern and Frankel, 2013, *et al*., 2017). Additionally, regulatory elements are developmentally dynamic, and consequently, their accessibility and activity can vary between tissue types and developmental timepoints. Delineating promoters and enhancers classes requires not only their actual genomic annotation, but also characterisation of their spatiotemporal activity and the transcription factor binding motifs, and sequence composition within each regulatory class.

Genome wide CRE annotation has relied primarily on high throughput sequencing based technologies such as ATAC-seq and ChIP-seq. These capture accessibility and histone modification profiles across broad genomic regions. Specifically H3K27ac is associated with active regulatory regions, widely used to support active enhancer annotation, and H3K4me3 which is enriched around active promoters and transcription start sites (Mora *et al*., 2015; Smith *et al*., 2023). In non-model systems comparable ATAC-seq and histone modification datasets are becoming increasingly available, with previous studies identifying broad characteristic patterns of variable regulatory elements across spatiotemporal conditions (e.g. Lewis *et al*., 2016; Lewis and Reed, 2018). However, distinguishing features associated with functionally distinct promoter and enhancer classes have yet to be fully resolved. Characterising differences in genomic localisation, spatiotemporal regulatory activity, and transcription-factor sequence composition between promoter and enhancer classes provides an opportunity to establish a more comprehensive view of the regulatory architecture in non-model organisms.

One such system are the *Heliconius* butterflies, which have emerged as a leading system for investigating the regulatory control of morphological variation driven by integration of comparative genomics, population genetics, and the mechanics of developmental biology (Joron *et al*., 2006) (Figure 1). Currently, genomic and functional studies have demonstrated that wing pattern variation is primarily driven by a small subset of major effect loci and associated genes with modular regulatory architectures (McMillan *et al*., 2020; Belleghem *et al,* 2021). Among these, *optix* and *cortex* represent canonical examples that determine broad pattern variation across Lepidoptera. Notably, phenotypic divergence at both loci is partially mediated via *cis*-regulatory variation rather than coding sequence evolution. For example, at the *optix* locus, comparative genomic analyses have revealed multiple non-coding regulatory regions implicated in wing patterning, demonstrating a modular regulatory architecture in which individual CREs drive discrete spatiotemporal domains of gene expression (Reed *et al*., 2011; Wallbank *et al., 2*016). Subsequent sequencing and CRISPR-Cas9 based experimentation functionally validated multiple *cis-*regulatory enhancers at the *optix* locus revealing enhancer activity is both modular and pleiotropic with regionally variable combinatorial sets of enhancers influencing spatiotemporal expression patterns across wing development (Lewis *et al., 2*019).

**Figure 1.**
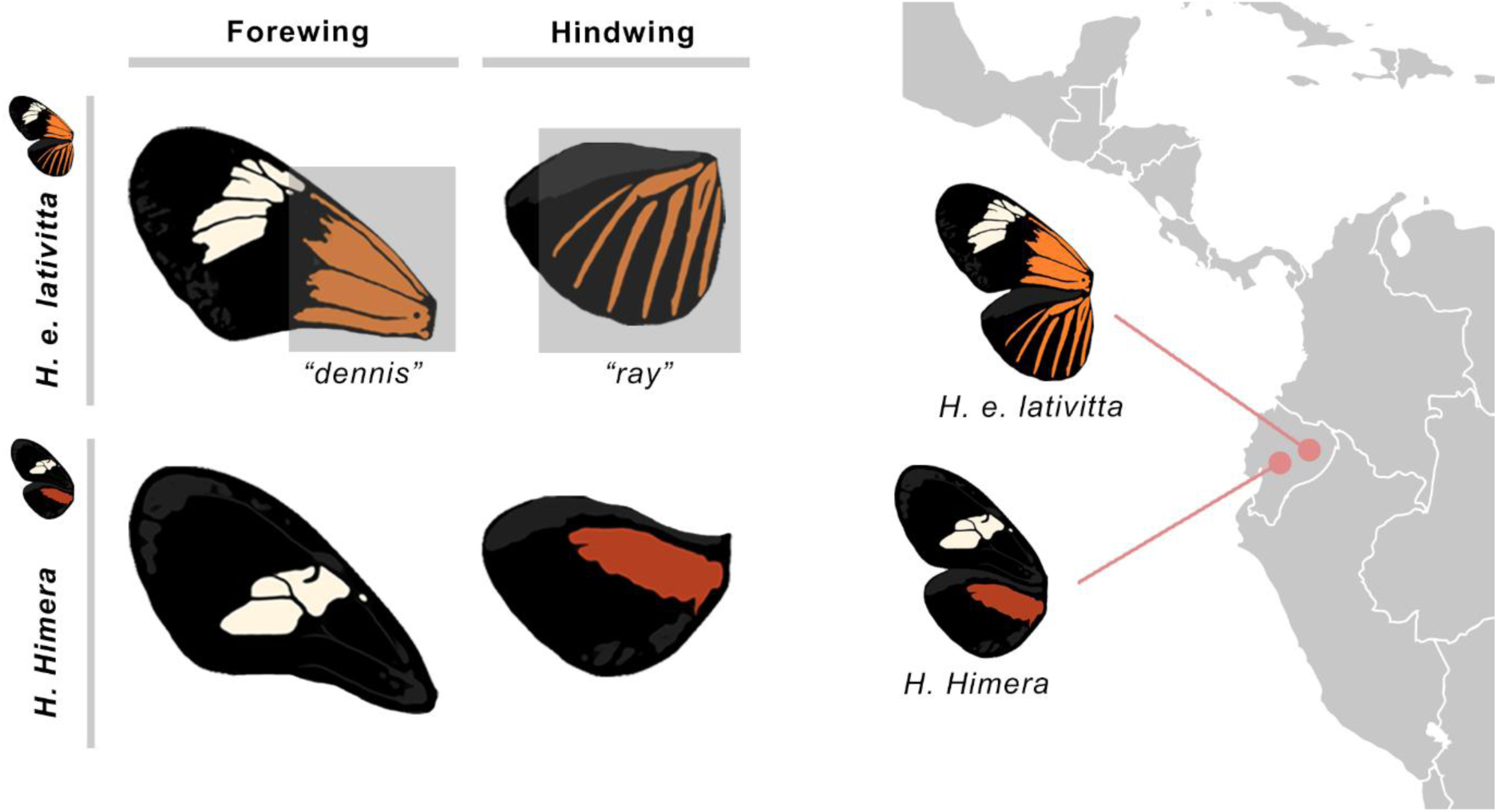
Overview of wing patterning diversification and geographic context of two *Heliconius* species: *Heliconius himera* and *Heliconius erato lativitta*.

*Cis*-regulatory activity is highly dynamic across *Heliconius* wing development, reflecting the temporal remodelling of regulatory programmes that first establish wing pattern identity and later drive scale differentiation and pigmentation. Specifically, in day 3, at the mid-pupal stage, wing tissue has reached its full size, and scale buds are beginning to develop, whereas the late ommochrome stage, approximately 7 days in, coincides with the development of visible red pigmentation associated with *optix* expression (Belleghem et al., 2021). Comparison of these stages therefore provides a framework for investigating temporal remodelling of regulatory activity during the transition from pattern establishment to pigmentation. Additionally, modular *cis-*regulatory organisation has been demonstrated broadly across the Lepidopteran taxa, with recent comparative analyses demonstrating that convergent wing pattern evolution is highly constrained with repeated use of conserved *cis-*regulatory architectures (Chehida *et al*., 2026).

While integrative epigenomic approaches have improved substantially for CRE annotation, with the rapid advancement of genome wide assays such as ATAC-seq and ChIP-seq, their application remains constrained by experimentally generated sequencing datasets which are currently available sporadically across varied spatiotemporal conditions for most non-model organisms, including *Heliconius* and related taxa. Consequently, machine learning approaches capable of regulatory element prediction directly from genomic sequences may extend CRE annotation beyond species with existing genomic resources. Machine learning methods have already provided a powerful framework for identifying complex, high dimensional patterns within genomic datasets that would otherwise require extensive empirical characterisation across a diverse range of vertebrates, insects, and plants to predict chromatin accessibility landscapes, enhancer activity, and functional non-coding regulatory regions (Kelley *et al*., 2016; Thibodeau *et al*., 2018; Sun *et al*., 2021; Kaplow *et al*., 2023; Wang *et al*., 2025). Establishing a baseline of regulatory features from experimentally characterised species therefore provides an essential foundation for developing predictive models capable of investigating CRE dynamics across a broader range of non-model organisms.

Convolutional neural networks trained via experimentally identified enhancer sequences provide a promising approach to expediting putative regulatory element annotation across species lacking equivalent epigenomic datasets. Additionally, independently available ATAC-seq data for a subset of *Heliconius* species provides an opportunity to assess whether predicted enhancer regions correspond to accessible chromatin outside species used for model training, thereby evaluating transferability of learned regulatory sequence architecture across species. Alongside this, comparative genomics provides an additional framework for investigating evolutionary conservation of regulatory regions. Specifically, conserved non-exonic elements (CNEEs) represent genomic regions which have evolved at slower rates than neutral sequences, therefore providing a method of identifying regions targeted by selection that have potential regulatory function (Edwards *et al*., 2017). Within Heliconiini, extensive comparative analysis has identified CNEEs with significant enrichment within accessible chromatin supporting their association with *cis*-regulatory function (Cicconardi *et al*., 2023; Cicconardi *et al*., 2025). Consequently, integration of CNEEs with machine learning predicted enhancers provides an opportunity to elucidate the extent of which putative regulatory regions are conserved across *Heliconius* and outgroup taxa.

Collectively, previous studies in *Heliconius* have shown that *cis-*regulatory architectures are crucial in elucidating the underlying mechanisms of wing patterning, and demonstrate that *cis-*regulatory variation can generate substantial phenotypic divergence while preserving protein-coding sequences (Reed *et al*., 2011; Wallbank *et al., 2*016; Lewis *et al., 2*019; Chehida *et al*., 2026). However, our current understanding of *cis-*regulatory mechanisms remains largely focused on a few major effect loci, and whilst individual CREs have been functionally validated around loci such as *optix*, broad characterisation of enhancer and promoter landscapes orchestrating the development of *Heliconius* wing and other tissues remain incomplete. This represents a crucial gap in our understanding. However, advances in high throughput chromatin accessibility, histone modification mapping, and extensive genomic data availability now enables systematic identification of a *cis-*regulatory atlas for *Heliconius* wing development (Cicconardi *et al*., 2023).

Here, we reanalyse publicly available ChIP and ATAC-seq datasets for multiple species of the *Heliconius* Erato clade. Substantial phenotypic variation in wing patterning is observed throughout the clade, exemplified here by *H. e. lativitta* and the closely related *H. himera*, which have been previous subjects of *cis-*regulatory characterisation (Lewis *et al*., 2019). Distributed throughout the western Amazon Basin, *H. e. lativitta* displays the Dennis–ray phenotype, consisting of a red proximal forewing patch and radiating red hindwing rays. In contrast, *H. himera* displays a yellow forewing band and red hindwing bar, occupying more arid, high-elevation habitats within Ecuador and Peru (Figure 1) (Supple *et al*., 2015; Lewis and Reed, 2018). Using data from these species, we delineate enhancer and promoter classes across *Heliconius* wing development characterising their genomic localisation, transcription factor association, and differential regulatory activity across spatiotemporal conditions in the two species. Additionally, we extend enhancer identification across the wider Heliconiini radiation using sequence-based machine learning, assess prediction transferability across species using independently available chromatin accessibility data, and finally integrate predicted enhancer regions with CNEEs to determine evolutionary conservation of putative regulatory regions.

## 2. Methods

### i. Data sets and preprocessing

Previously published ATAC-seq and ChIP-seq (H3K27ac and H3K4me3) datasets were obtained from the short read archive (SRA) (Lewis and Reed, 2018) for *H. himera* and *H. e. lativitta*. Sequence datasets consisted of forewing (FW) and hindwing (HW) tissues across two developmental timepoints, day 3 post-pupation (D3) and the ommochrome stage (OM; approximately 7 days post-pupation), with two biological replicates per condition. Full dataset information, including species, tissue, developmental timepoint, assay type, biological replicates and SRA accession identifiers, is provided in Supplementary Table S1, S2. Additional data was available for *H. e. petiverana*, however, technical limitations associated with regulatory element annotation prevented its inclusion in downstream comparative analyses (see Supplementary Materials Note).

Completeness and contiguity of reference assemblies for *H. himera* and *H. e. lativitta* was assessed using compleasm 0.2.10 (Huang and Li, 2023) and the lepidoptera odb12 database (Tegenfeldt *et al*., 2025). Sequence FastQ datasets for ChIP-seq (H3K27ac and H3K4me3) alongside ATAC-seq underwent the following data preprocessing pipeline. Adapter sequences and low-quality reads from ATAC-seq and ChIP-seq FastQ files were removed using trimgalore 0.6.10 followed by fastqc to assess and score the quality of trimmed reads. Subsequently multiqc was then used to create an aggregated fastqc report. Quality-filtered reads were then mapped to each species’ respective genome using bwa 0.7.17 (Li, 2013). Aligned reads were filtered to further remove low-quality reads with the parameter –q 20. Duplicate reads were removed from the filtered BAM files using picardtools 2.27.3 markduplicates (Broad Institute, 2018). Alignment quality, replicate concordance, and fragment size distributions were assessed using deeptools 3.5.5 (Ramirez *et al*., 2016).

### ii. Peak calling

To identify enriched ATAC-seq regions and ChIP-seq marks (H3K27ac and H3K4me3) peak calling was performed using macs 3.0.1 (Zhang *et al*., 2008). Peak calling parameters were specific to each sequencing type with additional input controls for ChIP-seq reads; ATAC: –f BAMPE –-nomodel--call-summit –B –q 0.05, H3K4me3: –-call-summit –B –q 0.05, and H3K27ac: –-broad –f BAMPE –B –q 0.01. Additionally, a custom python script SelectBestSummitMACS3narrowPeaks.py (available via GitHub repository in data availability section) then used to select the best peaks at overlapping locations based on the highest associated quality score for ATAC and H3K4me3 peaks. Subsequently, peak calling quality checks were then performed as follows: Fraction of Read in Peak (FRiP) scores were calculated for each sequencing type using samtools 1.3.1 (Danecek *et al*., 2021), and quality heatmaps were generated using deeptools. Additionally, the python script get_TSSandGeneBody.py (Rossello, 2024) and deeptools was used to visualise signal intensity 3kb around TSSs. Finally, TSS enrichment scores for ATAC data were generated via ataqv 1.3.1 (Orchard *et al*., 2020).

### ii. Characterisation of putative CREs

#### Identification

Species-specific putative CREs were identified independently per developmental stage, tissue, and replicate using bedtools2 2.31.1 intersect (Quinlan *et al*., 2010). Following established signatures of regulatory activity (Heintzman *et al*., 2007; Creyghton *et al*., 2010; Karchencko *et al*., 2011; Koenecke *et al*., 2016; Ivanković *et al*., 2024), promoter regions were defined as ATAC-seq peaks overlapping both H3K27ac and H3K4me3 peaks and located within ±1 kb of an annotated TSS, whilst active enhancers were defined as ATAC-seq peaks overlapping H3K27ac lacking H3K4me3 overlap and located outside ±1 kb of TSSs. For each spatiotemporal condition, CREs identified in either replicate were combined and merged using bedtools2 merge to generate a single promoter and enhancer dataset per species, tissue, and developmental stage. Permutation tests were used to assess likelihood of false identification of CREs using bedtools2 shuffle to create 10,000 replicates per dataset. Overall distributions of promoter and enhancer counts and signal intensity were plotted via a custom enrichmentCRE.R script (see data availability) Finally, signal intensity for CREs was plotted using deeptools.

#### Cis-regulatory mapping

Putative CRE peak regulatory locations were assessed using CisRegMap.py (see data availability). Regulatory regions were categorised into groups according to the following parameters on positive or minus orientation depending on the loci strand: exon and intron, derived from species-specific GFF3 gene annotations, with introns defined as gene-body regions excluding annotated exons; TSS-proximal, defined as 1000 bp upstream to +1000 bp downstream from the TSS; intermediate, defined as 1000 bp up to 5000 bp upstream the TSS; and distal, defined as any region outside these parameters (Karchencko *et al*., 2011; Koenecke *et al*., 2016). A randomised control set was generated using bedtools2 shuffle, creating 10,000 replicates per dataset from a shuffled replicate folder with an overlap fraction of 0.95.

#### Cis-regulatory association with orthologous genes

ATAC-seq and ChIP-seq signal profiles were examined around the candidate loci *optix* and *cortex.* Reference orthologs were mapped to *H. melpomene* from FlyBase. Protein FASTA files for *H. erato* subspecies were generated from genome annotations using gffread 0.12.8 (Pertea and Pertea, 2020). Reciprocal best hit orthologs were identified using diamond 2.1.7 (Buchfink *et al*., 2021) in combination with seqkit2 2.8.2 (Shen *et al*., 2024). Identified orthologs were then intersected with putative promoter and enhancer BED files using bedtools2. For enhancer assignment genomic intervals were extended ±250kb from gene bodies using bedtools2 slop to capture distal regulatory elements. Promoter regions were defined as the 2kb upstream of target gene TSSs and identified using bedtools2 window. Experimentally characterised enhancers for *optix* were obtained from previously published CRISPR sgRNAs (Lewis *et al*., 2019). SgRNAs were then aligned to the *H. e. lativitta* assembly using bwa aln and samse, and mapped enhancer fragments were then merged to generate BED intervals. Subsequently, halliftover was performed to match enhancer regions to *H. himera* (Zhang *et al*., 2020). Experimentally recovered enhancers were identified via intersection with the mapped enhancer intervals. Signal enrichment for ATAC-seq and ChIP-seq datasets alongside associated promoter and enhancer elements were visualised across ±250kb of the target gene TSS using bedtools2 slop and pygenometracks 3.9 (Lopez-Delisle *et al*., 2021).

### iii. Characterisation of CRE transcription factors

#### De novo and known motif analysis

Motif analysis was performed using homer 5.1 (Heinz *et al*., 2010). A background set for motif analysis was created using shuffled CRE BEDs and each species respective chromosome sizes. Motif analysis with homer’s findmotifsgenome.pl was conducted using parameters mset insect –size. Motifs were then clustered via rsat matrix-clustering_stand-alone (Castro Mondragon *et al., 2*017). Additionally, bedtools2 nuc and makewindows with parameter –w 1000 was used to calculate GC content per each reference assembly and CRE BED file prior to being plotted with a custom R script violinGCplot.R (see data availability).

#### Footprinting analysis

Footprinting analysis was performed using tobias 0.14.1 on merged putative CRE datasets for each species (Bentsen *et al*., 2020). ATAC-seq BAM files per species were merged via samtools consolidating all spatiotemporal and replicate conditions for each test species. From tobias atacorrect was applied to correct for Tn5 transposase bias. Subsequently tobias footprintscores was then used to calculate continuous footprinting scores across CRE regions. Homer motif analysis outputs were then converted from homer to meme format using homer2meme.py for compatibility with tobias bindetect to identify differential binding of transcription factors between *cis-*regulatory conditions in each species. Finally, tomtom 5.5.9 (Gupta *et al*., 2007) was used to identify motifs showing significant differential binding levels enhancer and promoters, transcription factors were then matched to the jaspar insects 2024 database (Rauluseviciute *et al*., 2024).

#### Differential expression of CREs across spatiotemporal conditions

Deseq2 was used to normalise read counts and determine differential expression of CREs across spatiotemporal conditions (Love *et al*., 2014). CRE BED files were first converted to simplified annotation format using AWK, and following this ATAC, H3K27ac, and H3K4me3 peaks pertaining to each CRE BED file were split into consensus sets. Counts for each consensus set was performed using featurecounts 2.1 (Liao *et al*., 2014), generating a matrix of raw counts per CRE across all samples. To reduce noise, peaks with low read support were filtered prior to differential analysis by removing features with a total count less than 10 across all samples. Quality control of sample replicate consistency was assessed using PCA of variance normalised counts to confirm expected clustering by biological condition. Additionally, MA plots were generated to visualise log2 fold-changes in promoter and enhancer chromatin and histone mark signatures across spatiotemporal conditions and assess global distribution of differential regulatory activity.

### iv. Prediction of CREs across Heliconiini genomes

#### Dataset construction

Sequence based enhancer classification was performed using experimentally defined *H. himera* and *H. e. lativitta* enhancers as positive examples and genomic regions not overlapping annotated enhancer regions as negative (background) examples. Enhancer intervals were converted to fixed 501bp windows centred on interval midpoints using bedtools2 slop, with incomplete scaffold edge windows removed. Sequence composition was calculated using bedtools2 nuc, and regions containing >1% ambiguous (N) bases were excluded. Candidate background regions were generated via partitioning of the genomes into non-overlapping 501bp windows using bedtools2 makewindows, and subsequently removing any annotated enhancers regions with bedtools2 intersect. Background regions were then sub sampled to match enhancer class weights using a custom python script matchbackground.py (see data availability section) which additionally matched the distribution of ambiguous N content in the enhancer datasets. Enhancer and background datasets were finally partitioned into training, validation, and test datasets at chromosome level using predefined chromosome splits consistent across species to prevent information leakage.

#### Model training

Genomic sequences corresponding to the training, validation, and test split windows were extracted from their respective species reference assemblies using grelu (Lal *et al*., 2025) intervals_to_strings and assigned binary background (0) or enhancer (1) labels. Initial model optimisation was performed using an optuna hyperparameter search on grelu’s dilatedconvmodel. To assess cross species generalisation, models were additionally trained on *H. himera* and evaluated via *H. e. lativitta* validation chromosomes which were withheld from training (‘held-out’ chromosomes), and reciprocally trained on *H. e. lativitta* and validated on *H. himera.* Following these assessments, enhancer and background sequences from both species were combined for final model training retaining identical chromosome level splits across the training, validation, and test partitions. Final model training was performed using the Adam optimiser (Kingma and Ba, 2014) with a learning rate of 8.61 x 10^−5^, batch size of 256, and a maximum of 100 epochs, with the best validation checkpoint retained for downstream predictions. Model performance was assessed on held-out validation and test datasets using area under the operating characteristic curve (AUROC, measures discrimination between positive and negative classes), with additional monitoring of area under the precision curve (AUPRC, summarises a classifiers precision-recall trade off), precision (proportion of predicted enhancers which are true enhancers), recall (proportion of true enhancers identified), F1 score (mean of precision and recall), specificity (proportion of background sequences correctly classified), and confusion matrix statistics (counts of true positives, true negatives, false positives, and false negatives).

#### Enhancer prediction

For genome wide prediction, the final model was applied across 34 Heliconiini genome assemblies, including *H. himera*, *H. e, lativitta*, and 32 additional assemblies described previously (Cicconardi *et al*., 2023) (Supplementary Figure S3). Assemblies were selected based on genome completeness and phylogenetic representation across the Heliconiini lineage, including representatives of *Eueides* and *Podotricha* alongside *Heliconius.* Each query assembly was partitioned into sliding 501bp windows at 100bp intervals using bedtools2 makewindows. Windows shorter than 501bp or containing non-ACGT bases were excluded, and remaining sequences were scored in batches using the final grelu model. Windows exceeding the model’s validation derived classification threshold were saved as predicted enhancer windows together with their respective model scores. TSS-proximal regions predictions were subsequently removed using bedtools2 intersect. Remaining positive windows were sorted and merged using bedtools2 merge to create contiguous predicted enhancer regions, retaining maximum prediction score and number of contributing windows for each merged enhancer interval.

Predicted enhancers were evaluated for enrichment in accessible chromatin regions using publicly available ATAC-seq datasets retrieved from NCBI Sequence Read Archive for *H. charithonia, H. e. demophoon,* and *H. melpomene* (Supplementary Table S3). Condition specific ATAC-seq peaks from forewing, hindwing, and head tissues were first analysed individually and then combined into species level union datasets using bedtools2 merge. Predicted enhancers were considered supported by ATAC-seq peaks where >50% of the ATAC interval overlapped a predicted enhancer regions using bedtools2 intersect –f 0.5. Enrichment relative to random genomic expectation was assessed using 1,000 permutations of the predicted enhancer datasets generated by bedtools2 shuffle, using –chrom to preserve chromosome assignment, *-*noOverlapping, and –excl to exclude regions within ±1kb of annotated TSSs. Observed ATAC enhancer overlap was compared with the permutation distribution to calculate fold enrichment and empirical p-values. Additionally, *cis*-regulatory mapping of predicted enhancer regions was performed via CisRegMap.py as described previously.

#### Downstream analysis

CNEE analysis was conducted using a previously curated *H. melpomene* CNEE dataset. CNEE coordinates were lifted from *H. melpomene* into each query species using halliftover –– nodupes against a 63-way species cactus whole genome alignment containing the 34 target species (Paten *et al*., 2011; Cicconardi *et al*., 2023). Overlap between CNEEs and predicted enhancer regions was assessed bidirectionally via bedtools intersect, calculating proportion of mapped CNEEs overlapping predicted enhancers, and vice versa. Liftover fragments sharing the same original CNEE identifier were counted as a single CNEE. Enrichment was assessed against 1,000 species-specific shuffled enhancer replicates, calculating permutation mean overlap, fold enrichment and empirical p-values for both overlap directions.

## Results

### i. Reference assembly and peak quality assessment

Genome completeness of the *H. e. lativitta* and *H. himera* reference assemblies was evaluated using compleasm analysis. *H. himera* presented the most complete genome at 97.6% whilst *H. e. lativitta*, scored 93.5%. Assembly continuity statistics included N50 and N90 scores which assessed scaffold or contig lengths above which 50% and 90% of the genome assembly is contained. *H. himera* exhibited the most contiguous assembly, with a scaffold N50 of 19.5Mb and N90 of 13.3Mb across a 408.5Mb genome. *H. e. lativitta* had a comparable assembly size of 418.4Mb, but substantially lower contiguity N50 and N90 scores of 5.48Mb and 1.43Mb, respectively, reflecting a more fragmented scaffold structure. Analysis of trimmed reads via fastqc presented results consistent with high-quality data preparation (Full multiqc report available on github). Assessment of replicate concordance displayed strong correlations between biological replicates within each condition and species (Supplementary Figure S1,S2). Notably however, forewing and hindwing tissues within the same replicate demonstrated higher correlation with each other than with the corresponding tissue type across different replicates, indicating limited tissue-specific divergence. Finally, ATAC library sizes were validated confirming a clear clustering periodicity of ∼50–200 bp in ATAC seq fragment length distributions confirming the expected fragment size distributions corresponding to nucleosome free and mononucleosome regions (Figure 2B).

**Figure 2.**
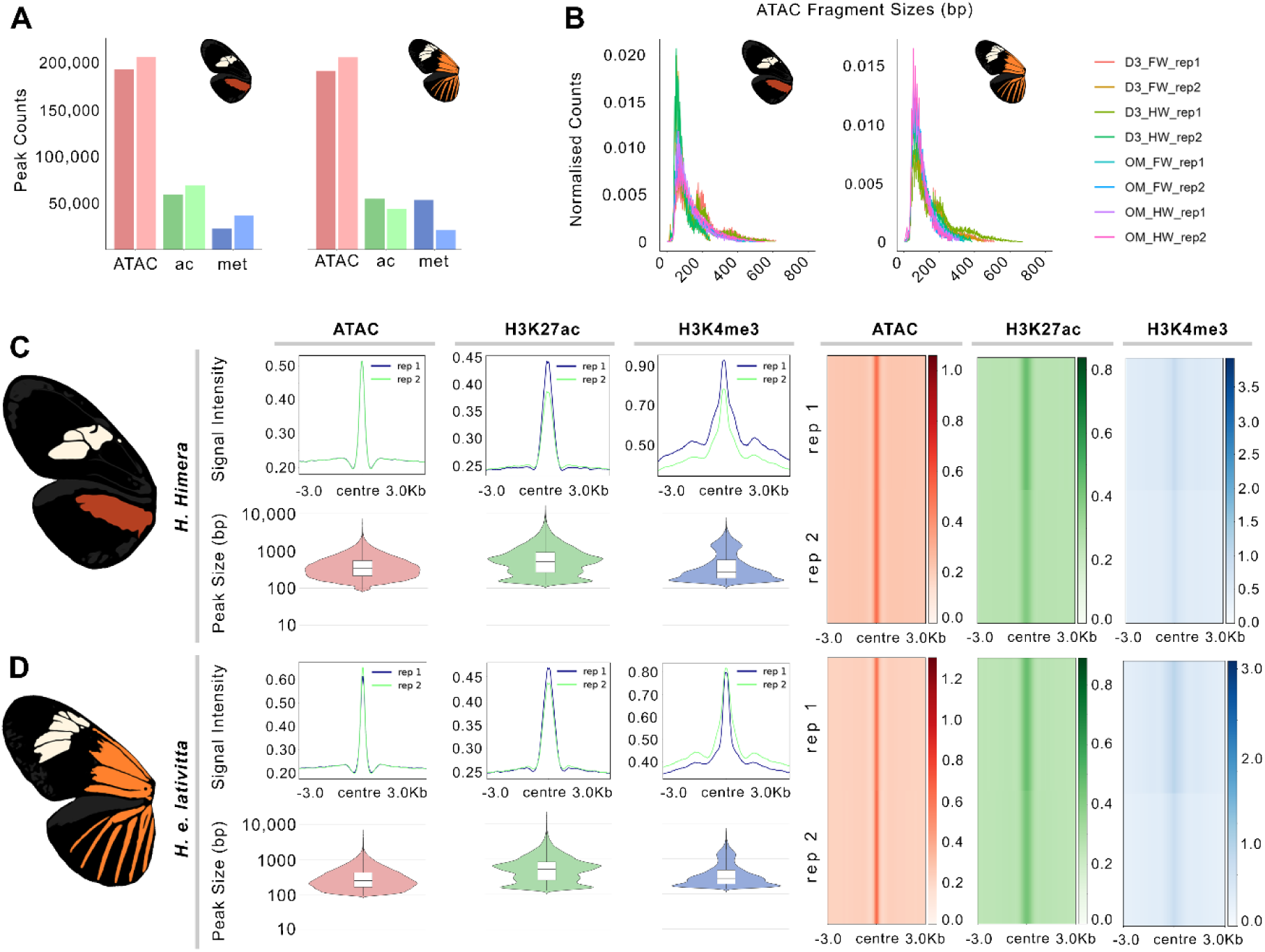
Preprocessing and peak calling quality checks for *Heliconius* data. (A) Overview of total peak counts for ATAC, H3K27ac, and H3K4me3 marks per species. (B) ATAC fragment size analysis for each species confirming high quality ATAC seq profiles. (C-D) Genome wide peak calling quality checks for both *Heliconius* species, showing coverage profiles in peak centred 3kb windows and heatmaps for each sequencing mark, alongside peak size distribution boxplots of all peaks used in downstream analysis.

Following macs3 peak calling merged ATAC-seq datasets contained 234,591 and 238,118 peaks, alongside 89,422 and 72,122 H3K27ac peaks, and finally 42,022 and 56,421 H3K4me3 peaks in *H. himera* and *H. e. lativitta* respectively (Figure 2A). FRiP scores were calculated per spatiotemporal condition, species, and replicate; ATAC-seq scores exceeded encode standards with the majority ranging from 0.45-0.60, with H3K27ac ranging from 0.07-0.23, and H3K4me3 ranging from 0.17-0.56. Low enrichment samples were evaluated for replicate concordance prior to inclusion in downstream universal peak set construction. Finally, ATAC TSS enrichment statistics were calculated, with *H. himera* averaging 4.06%, and *H. e.* lativitta at 3.36%. To assess signal enrichment for ATAC-seq and ChIP-seq data heatmap plots were generated revealing clear and consistent signal enrichment for peaks across all datasets (Figure 2C, D). In each dataset observed patterns of enriched signal intensity was characteristic of high-quality ATAC-seq and ChIP-seq that reflects precise localisation and reduced background noise in the corrected datasets.

### ii. Characterisation of putative promoters and enhancers

To identify and characterise CREs in *Heliconius*, datasets of putative promoters and enhancers were generated, leveraging the association of H3K27ac and H3K4me3 with enhancer and promoter activity. While it has previously been reported that these modifications may co-occur at enhancer and promoter elements, using a broad peak calling strategy for the H3K27ac mark alongside TSS based positioning mitigates overlap allowing for clear delineation of CRE classes, as demonstrated here and in supporting literature (Reed *et al*., 2016; Ivanković *et al*., 2024) (Figure 3).

**Figure 3.**
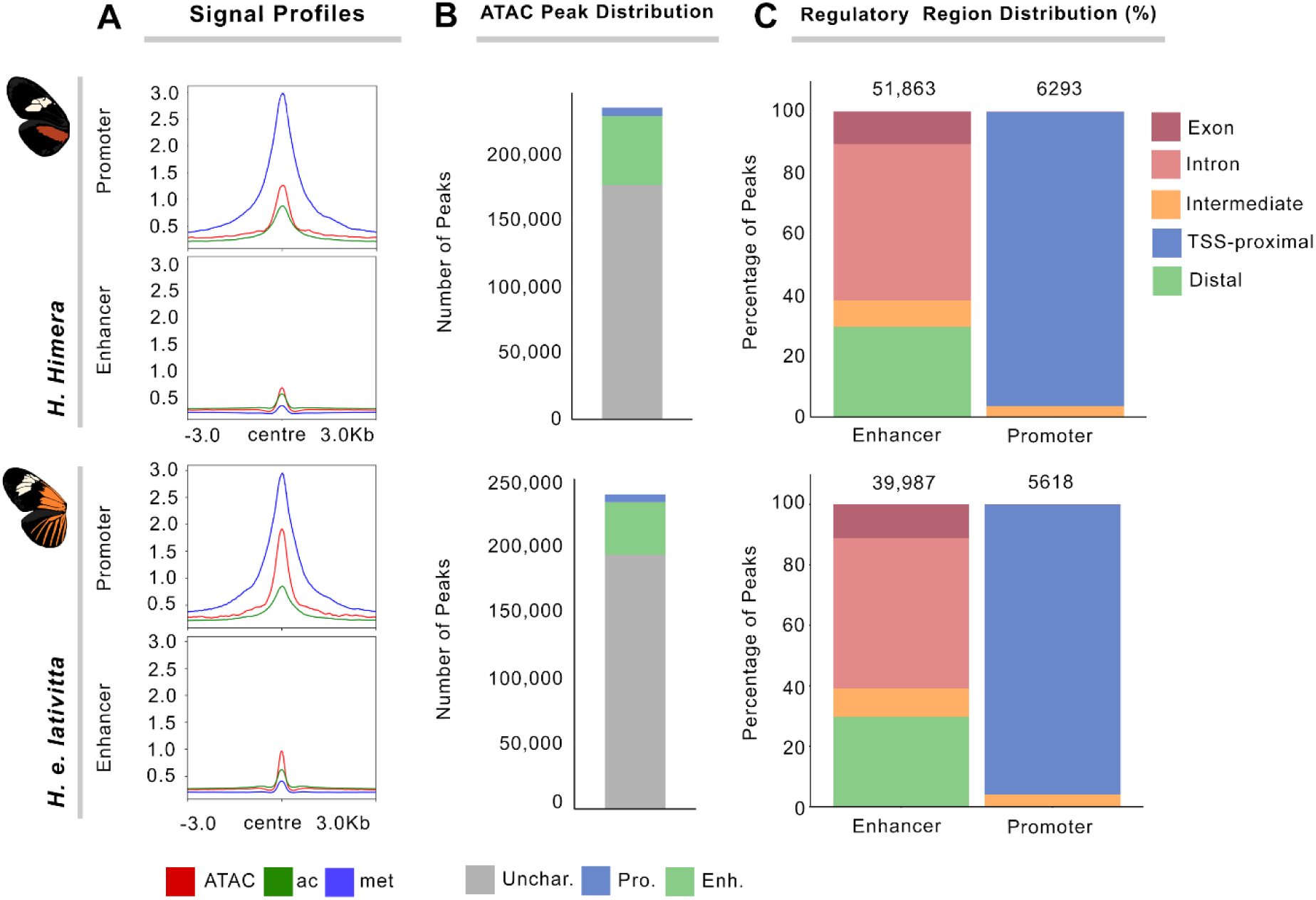
Promoter and enhancer dataset creation summary. (A) Displays signal profiles for ATAC, H3K27ac (ac), and H3K4me3 (met) present in each species promoter and enhancer BED files with signal intensity on y-axis and peaks 3kb around centres on x-axis. (B) Displays overall distribution of CREs across accessible chromatin landscape per species (%). (C) Displays *cis*-regulatory mapping plots for promoters and enhancers per species (%).

Following intersection, a total of 6,292 promoters and 51,862 enhancers were identified in *H. himera*, and 5,617 promoters and 39,986 enhancers were identified in *H. e. lativitta*. Promoter enhancer multiplicity in merged datasets was negligible, with 0.07% and 0.05% in *H. himera* and *H. e. lativitta* respectively. To evaluate significance of CRE occurrence against random chance, a permutation framework was applied in which ATAC peaks were randomly shuffled 1000 times to generate null distributions per promoter and enhancer condition. Enrichment scores were calculated as the ratio of observed to expected counts. Promoter and enhancer occurrence was enriched in both species, with fold enrichment of 1.45x/1.43x and 1.59x/1.76x for promoters/enhancers in *H. himera* and *H. e. lativitta,* respectively (p-value < 0.001 for all comparisons).

To characterise regulatory landscape distribution of putative CREs, promoter and enhancers *cis-*regulatory locations were analysed, quantifying the relative localisation of peaks across TSS-proximal, intermediate, intron, exon, and distal regions (Figure 3C). One-tailed binomial tests were conducted to compare observed number of peaks in each region to a randomised dataset to detect deviation from a random expectation. Promoter datasets displayed strong and consistent enrichment within TSS-proximal regions accounting for 85.6% and 83.6% of identified regions with significant overrepresentation detected in both *H. himera* and *H. e. lativitta* respectively (p < 0.01 for both comparisons). Conversely, gene bodies (3.7% and 3.5%), and distal regions (2.7% and 2.9%) were substantially underrepresented relative to random expectation within promoter datasets. (p < 0.01 for all comparisons). Enhancer datasets displayed broader genomic distributions and substantially weak TSS-proximal localisation (Figure 3C). In *H. himera*, enhancer peaks were strongly enriched within intronic regions, accounting for 51.2% of enhancers compared with an expected 38.8%, whilst exonic representation showed no significant deviation from random expectation (10.6% observed; 10.8% expected; p-value = 0.113). Distal and intermediate regions were modestly depleted relative to random expectation. Overall, enhancer localisation was considerably more distributed across different genomic localisations than promoters, with substantial representation within both intronic and distal sequences.

To determine whether putative CREs corresponded to known genes involved in *Heliconius* wing development, orthologs of *cortex* and *optix* were mapped to the two target *Heliconius* species from FlyBase and *H. melpomene*. Reciprocal best hit analysis identified high confidence orthologs of *cortex* and *optix* across the two *Heliconius* species from *H. melpomene* with low E-values and supporting high alignment scores (Figure 4). Overlap analysis between promoters and orthologs of *optix* and *cortex* revealed consistent promoter recovery across both *H. himera*, and *H. e. lativitta*. Analysis of distal enhancer overlap (±100 kb) with *optix* and *cortex* orthologs revealed higher levels of *optix* enhancer capture outnumbering *cortex* in both species suggestive of a more complex *cis-*regulatory architecture modulating *optix* expression. Recovery of experimentally characterised *optix* enhancers revealed of the five CRISPR validated elements, two overlapped the chromatin defined enhancer regions in *H. himera* (Figure 4A), and one in *H. e. lativitta* (Figure 4B). The remaining validated enhancer intervals did not overlap ATAC-seq peaks called under the criteria applied in our analysis and therefore did not meet the accessibility requirement established for enhancer classification.

**Figure 4.**
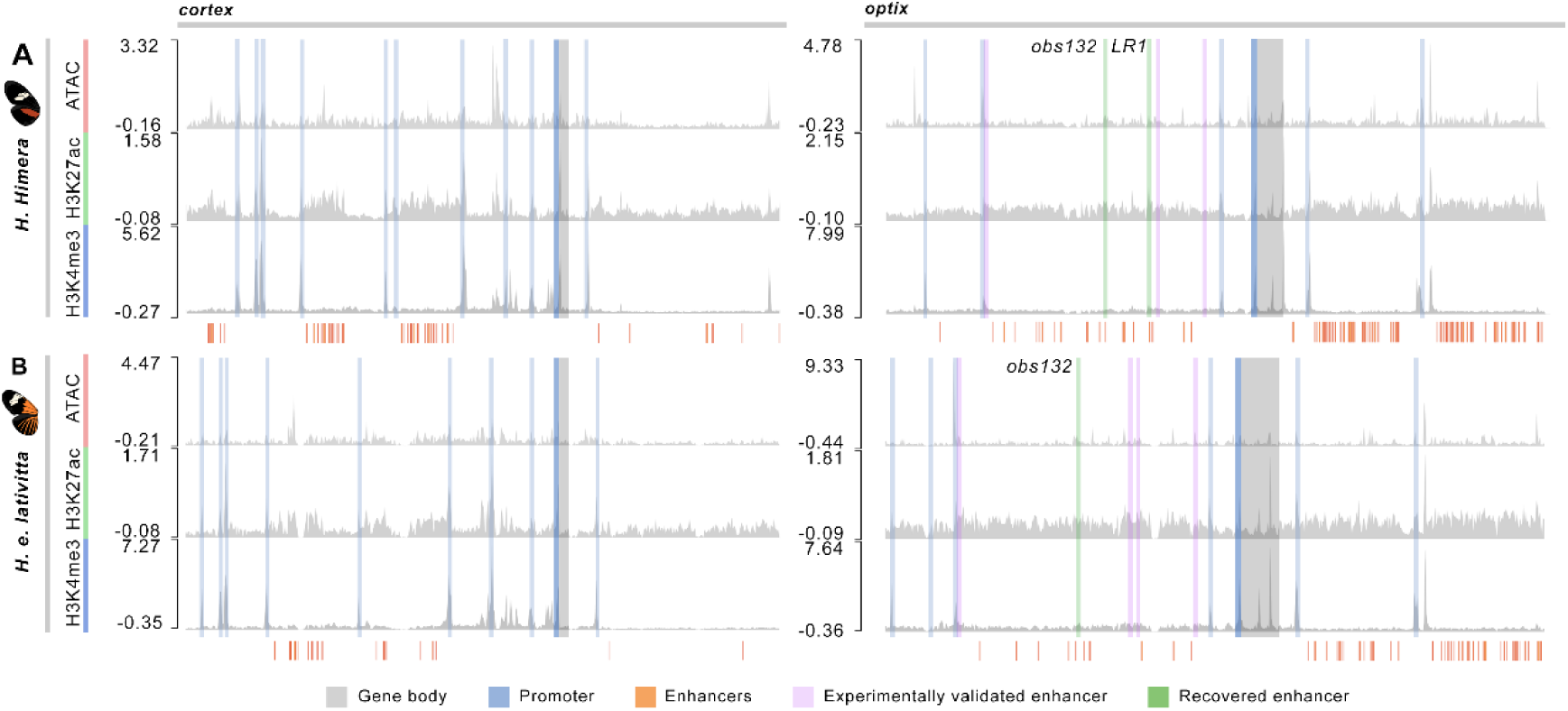
Signal track profiles in ∼250kb windows around the *cortex* and *optix* target genes including relative intensity scores for each signal track per gene and species. (A-B) Indicate successful capture of *cortex* and *optix* cognate promoters alongside additional CREs within the ∼250kb window. Recovery of the *obs132* enhancer of *optix* is additionally present in both species, whilst *LR1* is also recovered in *H. himera* marked in green, remaining experimentally validated enhancers from *optix* study presented in purple. All enhancers defined via histone and accessible chromatin overlap within the ∼250kb window are denoted by orange bars below each signal profile.

### iii. Characterisation of CRE transcription factors

To ascertain regulatory motifs within CRE datasets motif analysis was conducted on putative enhancer and promoter loci, allowing for identification of both known transcription factor binding motifs and overrepresented *de novo* sequence patterns, providing insight into the regulatory landscape of open-chromatin regions and the sequence level transcription architecture underlying enhancer and promoter activity. Additionally, GC bias within each of the CRE datasets and their respective reference assemblies was analysed (Figure 5B, D). Across both species, CRE datasets exhibited increased GC content relative to their genomic background. Genome wide GC content was consistent across assemblies with means of ∼0.31-0.33. Promoters showed moderate GC enrichment exhibiting mean GC contents ranging from ∼0.36-0.38 across species, while enhancers consistently displayed the highest GC content with mean values of ∼0.40-0.41 representing an approximate +0.07-0.10 increase relative to genomic background. (Figure 5).

**Figure 5.**
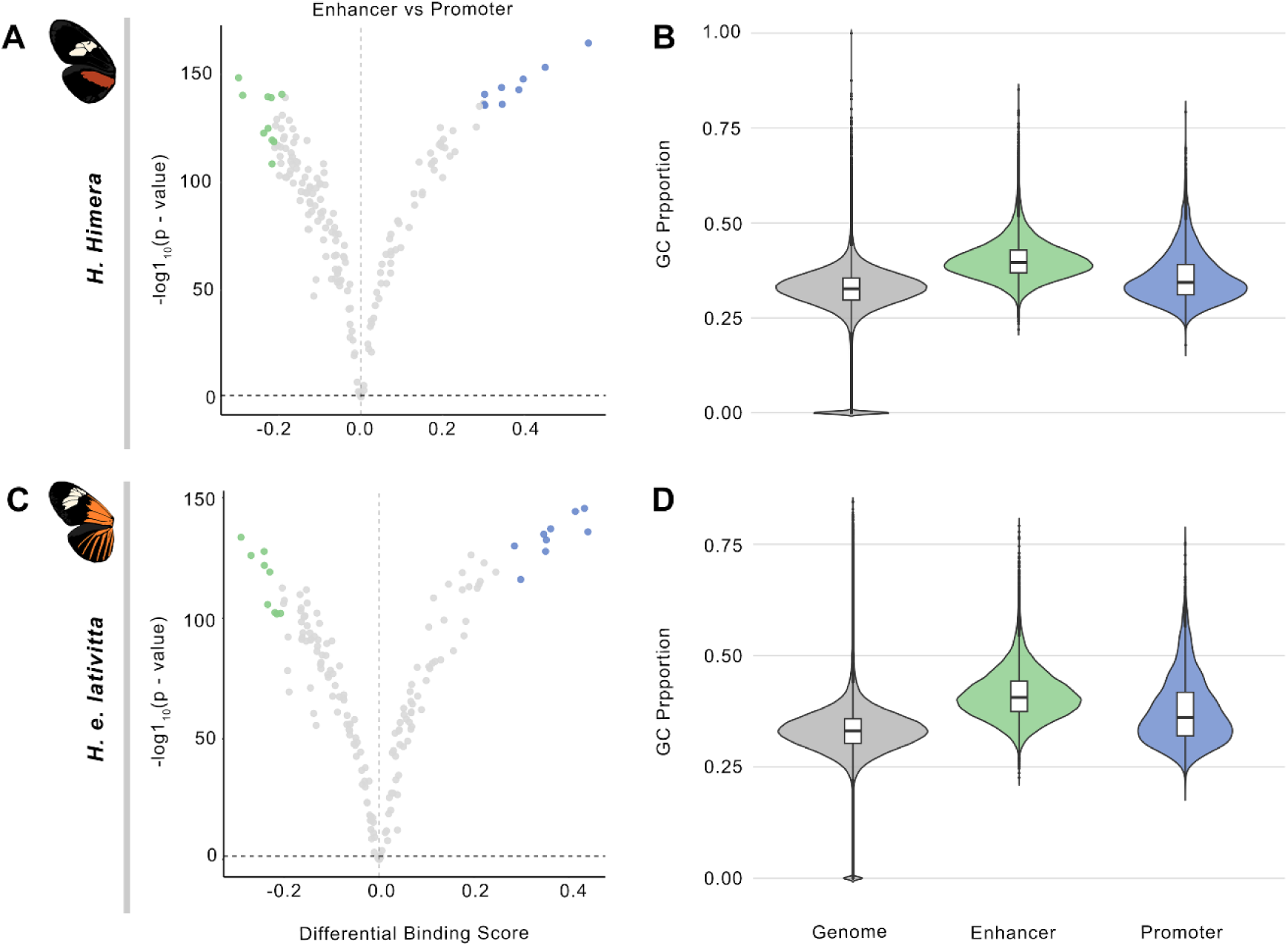
Displays summary of differential motif binding analysis between CRE datasets alongside GC enrichment. (A, C) Displays differential log2 fold change binding scores on x-axis of motifs identified in promoters and enhancer per species with negative log of p-value on y-axis, indicating shift towards conserved high affinity binding motifs in promoters and more densely clustered distribution of enhancer motifs in H. himera and H. e. lativitta. (B,D) GC content (%) of genome wide background, enhancer, and promoter regions in H. himera and H. e. lativitta. Significant differences were observed between all three categories in both species (Kruskal-Wallis, p-value < 2.2 x 10^−16^; pairwise Wilcoxon rank-sum tests with Benjamini-Hochberg correction, adjusted p-value < 2 x 10^−16^ for all comparisons).

Motif clustering revealed that enhancers exhibited higher motif diversity with cluster:motif ratios of 0.81 in *H. himera* and 0.77 in *H. e. lativitta*. While promoters exhibited ratios of 0.56 and 0.66 respectively. Across both promoter and enhancer datasets the majority of these clusters were single family motifs, however greater motif heterogeneity across both species indicated higher reuse of related regulatory factors within promoters.

To ascertain specifics of transcription factor (TF) binding, footprinting was carried out on identified motifs via the TOBIAS toolkit. Single run BINDetect on putative promoters and enhancers within each of both species enabled elucidation of differential binding between *cis-*regulatory TFs. Highlighted datapoints were measured as the top 5% of up-regulated and down-regulated differential binding scores (Figure 5A, C). Differential TF binding analysis displayed promoter specific binding signatures with motifs showing larger effect sizes of ∼0.45-0.60 in comparison to enhancers at approximately ∼0.25-0.30. Additionally, across the remaining datapoints there was a higher proportion of motifs displaying enhancer biased binding signatures, comprising 65.4% of non-highlighted motifs in *H. himera* and 57.3% in *H. e. lativitta*. This pattern was consistent with the slight overall negative shift in differential binding scores observed in both species, suggesting broader enhancer motif diversity whilst promoters contained stronger differential binding signals, coinciding with the previous clustering results.

To identify specific TFs and their associated genes, motifs showing statistically significant differential binding in promoters or enhancers within each species were matched to the JASPAR insects 2024 motif database via TOMTOM at a q-value confidence level of 0.1. In total, 24 TF matches were identified across *H. himera* and *H. e. lativitta*. For promoters 11 TFs were identified, nine of which were at a q-value < 0.05, six of these < 0.01 indicating several high confidence matches. Additionally, 13 enhancer matches were identified from the dataset, with 12 passing the q-value threshold of 0.05, and eight high confidence matches < 0.01. Nearly all identified TFs have been noted as displaying low to very high expression in *Drosophila melanogaster* wing discs suggestive of similar expression in *Heliconius* wing development. The majority of these have additionally been implicated in developmental processes, including myogenesis, neurogenesis, organ development, cell fate specification, and wing development. Enhancer associated matches were largely represented by basic helix-loop-helix and related transcription factor families, alongside factors involved in Notch and ecdysone signalling, suggesting enhancer regulation may be associated with developmental signalling and temporally dynamic transcriptional control. Several matches in *H. e. lativitta* additionally were members of the *Enhancer of split* complex.

### iv. Differential accessibility of CREs across spatiotemporal conditions

Differences in CRE accessibility was characterised across tissue types and developmental timepoints using signal intensity of ATAC, H3K27ac, and H3K4me3 marks associated with putative promoters and enhancers. Low differential binding scores relating to chromatin accessibility and histone mark enrichment highlights regulatory elements which exhibit stable accessibility levels across spatiotemporal conditions. In contrast, variation in ATAC signal reflecting dynamic changes in chromatin accessibility alongside differential enrichment of H3K27ac and H3K4me3 marks which distinguish dynamic activation of enhancers and promoters, highlighting CREs which may contribute to context-specific phenotypic divergence across spatiotemporal conditions in *Heliconius* wing patterning (Figure 6).

**Figure 6.**
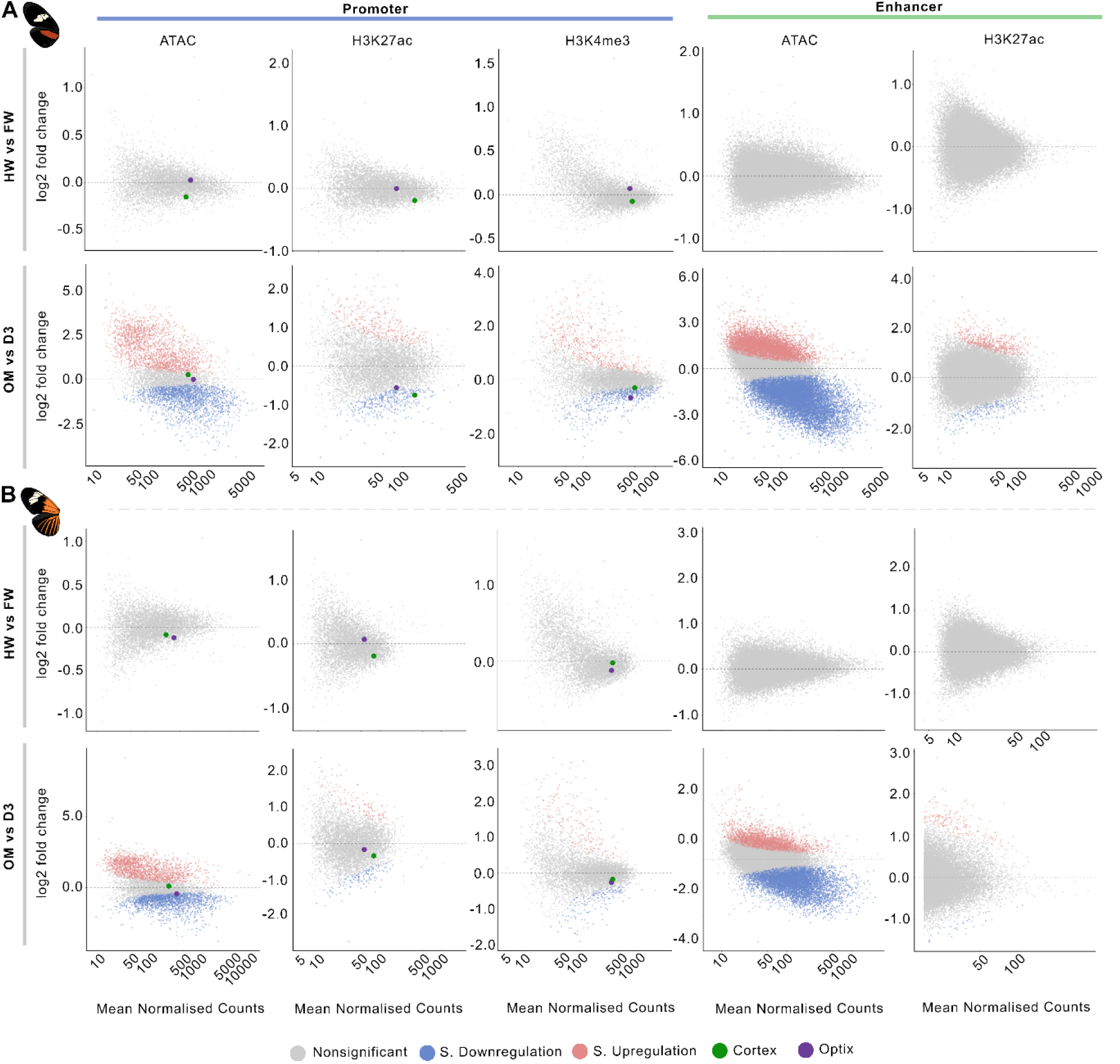
Displays differential enrichment analysis of signal types corresponding to promoters and enhancer across spatiotemporal conditions with log2 fold-change displayed on y-axis and mean normalised counts per signal type on y-axis. (A-B) *H. himera* and *H. e. lativitta* display highly conserved trends of minimal differential expression between hindwing and forewing tissue types, with considerably higher dynamicity between temporal development. Promoters at TSS of *cortex* and *optix* are additionally highlighted, showing minimal difference in accessibility or histone enrichment across tissue types, however statistical significance in *H. himera* histone enrichment across developmental timepoints.

Comparative analysis of CRE chromatin accessibility and histone mark enrichment displayed a high degree of conservation in CRE activity between *Heliconius* forewing and hindwing tissue types. In both promoter and enhancer datasets, very few peaks were found to display statistically significant differential accessibility with signal distributions clustered around zero log2 fold-change indicating minimal tissue specific changes. This pattern was consistent across ATAC, H3K27ac, and H3K4me3 marks in *H. himera* (Figure 6A) and *H. e. lativitta* (Figure 6B) supporting extensive reuse of regulatory elements across wing regions. In addition to genome wide trends, *optix* and *cortex* specific promoter analysis further supported the apparent lack of tissue specific divergence. In *H. himera* and *H. e. lativitta*, both *cortex* and *optix* associated promoters exhibited no significant differential enrichment with minimal change across ATAC, H3K27ac, and H3K4me3 datasets and log2 fold-change values consistently close to zero (log2FC < 0.3 across signal types) and high adjusted p-values (p value < 0.05), reinforcing tissue specific promoter activity is highly stable at these loci.

Differential chromatin accessibility and histone marks between developmental timepoints revealed widespread divergence in CRE regulatory states as compared to the conserved tissue profiles. Promoter associated accessibility displayed a greater prevalence of significant change in both species, with 68.8% of promoters and 54.4% of enhancers in *H. himera* showing significant differential ATAC signal (adjusted p-value < 0.05) with corresponding 62.2% and 34.9% values in *H. e. lativitta*. However, constraining statistically significant differential values to absolute log2 fold-change ≥1, difference between regulatory classes was substantially reduced. In *H. himera*, 41.5% of promoters and 38.3% of enhancers exceeded absolute log2 fold-change ≥1, and 32.4% of promoters and 23.8% of enhancers in *H. e. lativitta*. Consequently, whilst promoter regions showed a greater overall prevalence of statistically significant differential accessibility, a substantial proportion of these changes were comparatively moderate in size.

Histone modification profiles were comparatively more conserved between developmental timepoints than chromatin accessibility. Differential H3K4me3 promoter enrichment was measured at 20.2% of *H. himera*, and 6.7% of *H. e. lativitta* whilst H3K27ac enrichment different significantly across 12.2% of promoters and only 1.6% of enhancers in *H. himera*, with corresponding values of 5.1% of promoters and 1.3% of enhancers in *H. e. lativitta*. When additionally requiring log2 fold-change of > 1, these values decreased to 5.1% of promoters and 1.5% of enhancers in *H. himera,* and 2.4% and 1.0% in *H. e lativitta* respectively. Collectively, these results supported substantial chromatin accessibility remodelling across developmental timepoints, whilst histone modification profiles showed comparatively greater levels of conservation particularly within H3K27ac enrichment across enhancers.

### v. Sequence based enhancer prediction and CNEE conservation

Following sequence and *N*-content filtering, 51,268 *H. himera* enhancer sequences and 39,206 *H. e. lativitta* enhancer sequences were retained. Background selection comprised content distribution consistent with the enhancer datasets with mean *N*-fractions of 0.5×10^−6^ and 5.99×10^−4^ for *H. himera* and *H. e. lativitta* respectively. Reciprocal cross species evaluation demonstrated consistent sequence-based enhancer classification between the two species. Models trained on *H. himera* and evaluated on held-out *H. e. lativitta* datasets achieved test values of 0.894 AUROC and 0.904 AUPRC, whilst models trained on *H. e. lativitta* and evaluated on held-out *H. himera* datasets achieved values of 0.909 and 0.908 AUROC and AUPRC respectively. At validation derived thresholds corresponding approximately to 90% precision, test precision was 0.907 and 0.918 with a recall of 0.631 and 0.531 respectively demonstrating considerable cross species generalisation for enhancer prediction. Following reciprocal evaluation, sequences from both species were combined for final model training.

Combined model performance achieved validation AUROC and AUPRC values of 0.906 and 0.901, increasing to 0.909 and 0.913 on the independent test dataset. Validation derived classification threshold of 0.798 displayed test precision of 0.912 and recall of 0.625, specificity of 0.923 and an F1 score of 0.741. Mean prediction scores displayed clear delineation of enhancer and background sequences in the test dataset at 0.783 and 0.259 respectively, supporting consistent discrimination of regulatory and background sequences. The sequence model was subsequently used to predict enhancer regions across 34 Heliconiini reference assemblies, including both *H. himera* and *H. e. lativitta*. Within the experimentally characterised species, in *H. himera* the model recovered 73.3% of experimental enhancers at >1bp overlap, and 69.7% when requiring >50% of the experimental enhancer to be overlapped. For *H. e. lativitta*, recovery is 65.2% at >1bp and 62.0% at >50%bp. The close correspondence between 1bp and 50% recovery values indicates that majority of the recovered experimental enhancers cover a substantial region of the experimental interval rather than recovery percentage being bolstered by marginal overlap at interval boundaries. At the locus resolution, sequence-based predictions also showed strong correspondence with experimentally validated enhancers at *optix*, outperforming our previous analysis with ATAC and histone mark data. Of the five CRISPR-validated enhancers, four were recovered in both *H. himera* and *H. e. lativitta*, providing further support for the sequence prediction models capture of genuine regulatory signal (Figure 7A, B).

**Figure 7.**
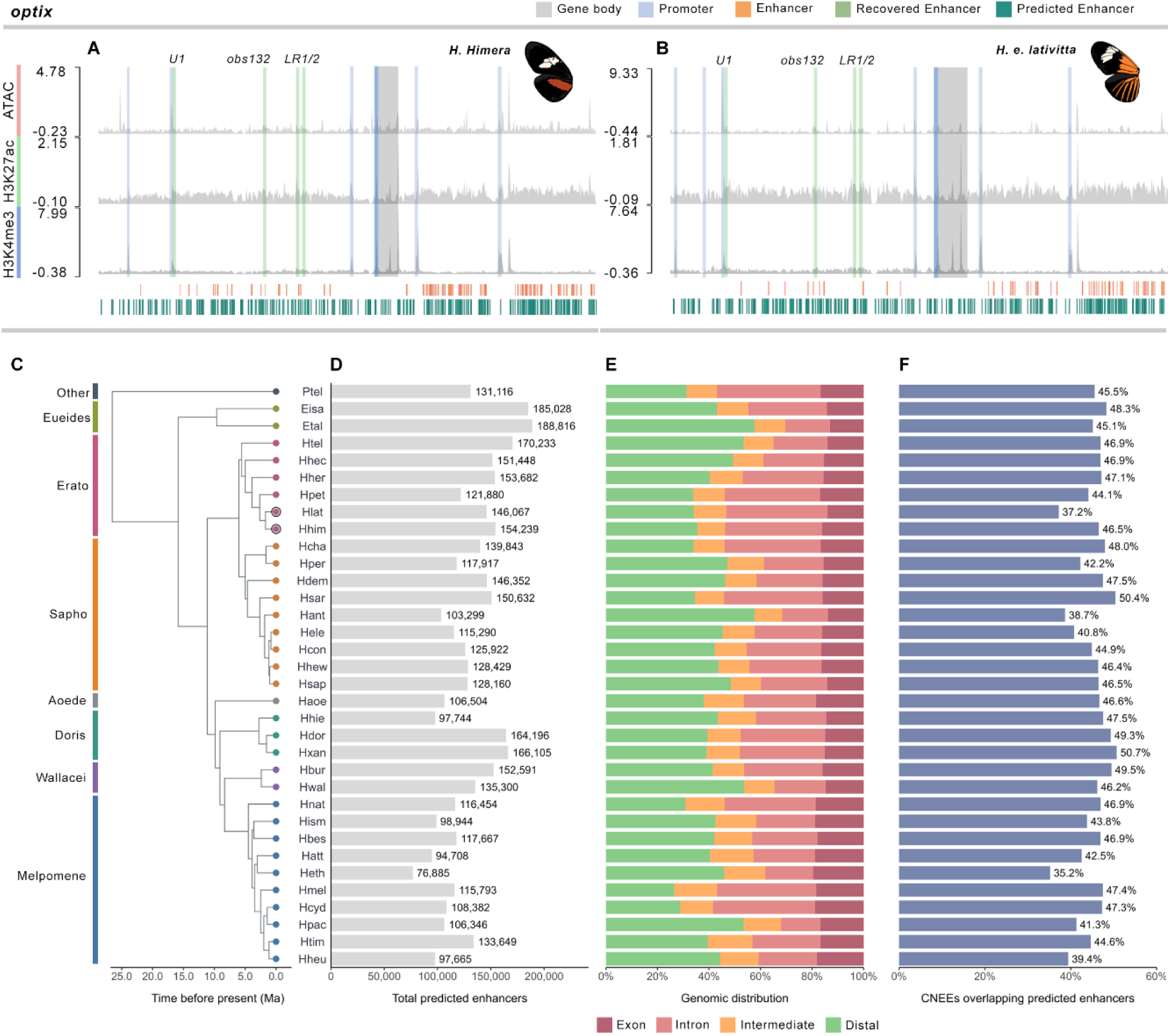
Overview of enhancer prediction analysis. (A-B) Displayed 250kb windows around *optix* for *H. himera* and *H. e. lativitta*, including predicted enhancer tracks demonstrating higher recall of 4/5 experimentally characterised enhancers, surpassing recovery demonstrated by ATAC and histone-based identification. Predicted and experimentally defined enhancers are displayed with individual bars under each signal track profile. (C) Phylogenetic context for the selected Heliconiini species with training species *H. himera* and *H. e. lativitta* highlighted with black ring (D) Total predicted enhancer counts across all 34 test species (mean = 130,577: IQR = 111,836-151,040). (E) Regulatory mapping for predicted enhancer regions, showing highest enrichment in distal and intron regions). (F) Overlap percentages of CNEEs within predicted enhancer regions.

To assess cross species generalisability of the sequence model, enhancer predictions were intersected with independently available ATAC-seq datasets from species representing three phylogenetically distinct lineages within *Heliconius*, *H. melpomene, H. charithonia,* and *H. e. demophoon* from the Melpomene, Sara/Sapho, and Erato clades respectively (Kozak *et al*., 2015), reflecting the full phylogenetic breadth of the *Heliconius* genus. Across merged ATAC datasets, predicted enhancers overlapped 54.6% of accessible regions in *H. melpomene,* 50.8% in *H. charithonia*, and 50.2% in *H. e. demophoon*. Notably, despite enhancer training data pertaining to wing tissues, substantial overlap of predicted enhancers was also observed within head ATAC-seq data, reaching 59.1%, 53.4%, and 53.7% in *H. melpomene, H. charithonia,* and *H. e. demophoon* respectively. Permutation analysis indicated that enrichment was substantially greater than random expectation across the genome, with values of 2.09x, 2.09x, and 1.86x fold enrichment above shuffled datasets for *H. melpomene, H. charithonia*, and *H. e. demophoon* respectively (p-value < 0.001). Overall, the sequence prediction model displayed transferability across the *Heliconius* clade, while substantial correspondence with head chromatin accessibility suggests that the learned regulatory sequence characteristics are not restricted to wing tissues.

To determine enrichment of predicted enhancers corresponding to evolutionarily conserved regulatory sequences across Heliconiini, enhancer datasets were intersected with CNEE regions. Substantial overlap was detected across all 34 species, with 35.2-50.7% of CNEEs intersecting predicted enhancer regions (mean 45.3%) (Figure 7F). Reciprocal analysis revealed 22.3-39.3% of predicted enhancer regions overlapping CNEEs (mean 32.3%), indicating approximately one third of predicted enhancer regions contained evolutionary conserved non coding sequences. CNEE overlap by predicted enhancer regions maintained broadly similar values across major Heliconiini lineages, with mean overlap ranging from 44-49% suggesting limited clade specific variation in the conservation of non-exonic elements captured by the model. Fold enrichment demonstrated that overlap was greater than random expectation across all species in both directions. CNEE-to-enhancer enrichment typically ranged from 1.61–1.71 fold, while reciprocal enhancer-to-CNEE enrichment ranged from 1.25–1.34 fold. Additionally, *H. himera* displayed the highest overlap in both directions, at 1.92x and 1.52x respectively, whilst *Heliconius pachinus* displayed the lowest values at 1.45x and 1.18x.

## Discussion

Through the analysis of *Heliconius cis-*regulatory architecture we aimed to provide the first large scale delineation and characterisation of *cis-*regulatory promoters and enhancers in *Heliconius*, and to elucidate CRE dynamics involved in *Heliconius* wing development across spatiotemporal conditions. By integrating analysis of chromatin accessibility through ATAC seq and histone modifications via the H3K27ac and H3K4me3 marks, high confidence peak sets following stringent quality checks allowed the annotation of putative promoters and enhancers which displayed genomic distribution consistent with canonical regulatory organisation (Ivanković *et al*., 2024). Annotation of CREs integrating chromatin accessibility and histone signatures is inherently complicated due to overlap between enhancer and promoter associated marks, as previously noted (Lewis *et al*., 2016). Optimisation of the peak calling strategy mitigated this ambiguity in delineating promoters and enhancers via switching to a broad peak calling system for the H3K27ac mark and applying TSS based definition, providing a framework for interrogating distinct regulatory classes. Following additional TSS-based classification, promoters were concentrated within TSS-proximal regions, while enhancers displayed a broader genomic distribution across gene body, intermediate, and distal regulatory regions, consistent with their established long-range and context-dependent functions (Pennacchio *et al*., 2013).

In *H. melpomene* there are approximately ∼13,000 annotated genes with expression breadth for other lepidopteran species reporting ∼30-60% are actively transcribed during specific spatiotemporal conditions (Dasmahapatra *et al*., 2012; Cao and Jiang, 2017). Consequently the 6,460 *H. himera*, and 4,553 *H. e. lativitta* promoters appear to fall within the expected range for actively transcribed loci. Similarly, promoter and enhancer association at the *optix* and *cortex* provide further support for successful capture of putative CREs. Distal enhancer association were more prevalent than promoters, particularly at the *optix* locus for both species. This is consistent with previous work demonstrating that *optix* is characterised by pleiotropic effects on wing patterning and venation regulated by combinatorial action of numerous enhancers (Lewis *et al*., 2019). Extension of these experimentally defined enhancer sequence characteristics through the cross-species prediction model, with predicted enhancers displaying substantial enrichment of accessible chromatin and association with CNEEs across the wider Heliconiini phylogeny. Collectively, these results provide a comprehensive annotation of promoters and enhancers, alongside those associated with key wing patterning genes *optix* and *cortex* providing a CRE landscape for functional stratification of sequence architecture, regulatory grammar, and developmental dynamics underlying phenotypic diversification, whilst additionally extending enhancer sequence characterisation and evolutionary conservation across the wider Heliconiini phylogeny.

### i. Sequence and motif architecture distinguish regulatory classes

Sequence composition and motif architecture provides support for functional differentiation between putative promoters and enhancers. Across both species, CREs exhibited higher GC contents relative to their genomic background consistent with enrichment for transcription factor binding sites (Wang *et al*., 2012). Notably, GC enrichment was highest in enhancer datasets coinciding with increased motif diversity, this contrasts with core vertebrate promoters which are typically associated with higher GC contents due to enrichment of CpG islands. However, CpG island enrichment in promoters has not been corroborated in invertebrates such as *D. melanogaster* which may explain the minor increase in promoter GC content compared to genomic background (Kowal *et al*., 2020). The patterns observed here are consistent with broader observations in insect regulatory systems, where promoter associated CpG island enrichment is less pronounced than in vertebrates (Kowal *et al*., 2020). In Hymenoptera, for example, CpG methylation primarily occurs in clustered gene body regions rather than promoters, indicating a fundamentally different genomic organisation of CpG associated regulatory features compared to vertebrates (Jeong *et al*., 2018). Consequently, GC rich *cis-*regulatory regions in *Heliconius*, like other insect systems, may be associated with increased transcription factor binding site density reflecting their role in modulating gene expression via combinatorial transcription control.

Motif clustering analysis further supports distinction in *cis-*regulatory architecture, with enhancer datasets consistently exhibiting higher motif diversity in comparison to promoters. Motif clustering patterns observed in this study are consistent with previous work displaying enhancer regions characterised by high motif diversity and flexible transcription factor association whilst promoters exhibit a more conserved regulatory architecture with greater motif clustering of core transcriptional machinery (Yang *et al*., 2017; Zhu and Landsman, 43 2023). Elevated motif diversity identified within enhancers may be reflected in *Heliconius* wing patterning development, where precise spatiotemporal regulation requires the integration of multiple developmental and tissue specific transcription factors. This is consistent with previous characterisation of key wing patterning genes such as *optix*, which is modulated by multiple distinct enhancers driving context specific gene expression, illustrating how modular enhancer organisation enables precise control of phenotypic diversity (Lewis *et al*., 2019. Recent analyses have found in vertebrate systems TF binding site composition between promoters and enhancers may partially reflect CpG island associated sequences rather than fundamentally distinct regulatory architectures (Andersson *et al*., 2014). However, as noted, CpG island enrichment at CREs is substantially reduced in invertebrate genomes including insects, consequently the observed motif divergence in *Heliconius* CREs may reflect genuine divergence in transcription factor binding site organisation between promoters and enhancers.

### ii. Differential transcription factor binding reveals distinct regulatory logic

Differential binding patterns between CREs reinforced distinct regulatory logic between putative promoter and enhancer datasets. Promoter associated motifs displayed greater differential binding indicating fewer but stronger signal TF interactions than enhancers which displayed reduced individual differential binding scores though higher numbers of differentially bound motifs consistent with broader TF utilisation and combinatorial binding which coincides with previously reported trends between CREs (Nguyen *et al*., 2016; Reiter *et al*., 2017; Lewis *et al*., 2019). Identification of 24 TFs linked to roles in developmental processes including wing, neural, muscular, and cell fate development indicates functional regulatory activity within the characterised CRE datasets. Enhancer associated motifs were particularly represented by basic helix-loop-helix and related transcription factor families alongside factors associated with Notch and ecdysone signalling, consistent with more context-dependent and developmentally responsive regulation. Overall differential binding trends observed conform to established models of more conserved promoter regulatory architecture, whilst enhancers provide greater flexibility and context-dependent control through combinatorial binding and broader transcription factor repertoires (Nguyen *et al*., 2016; Rao *et al*., 2022).

### iii. CRE activity stable across tissues but dynamic across development

Accessibility of chromatin and histone mark enrichment maintained stability in both promoters and enhancers across tissue types, suggesting minimal CRE dynamicity involved in differential phenotypic development between hindwing and forewing tissues. Slight differences, though none statistically significant, were noted in *optix* and *cortex* promoter differential chromatin accessibility in *H. himera* and *H. e. lativitta* correlating with their respective pigmentation phenotypes, i.e. higher accessibility of both hindwing *optix* promoter compared with forewing involved in red pigmentation, and the forewing *cortex* promoter involved in yellow/white pigmentation in *H. himera* (McMillan *et al*., 2020). Whilst differential expression of CREs appears principally responsible for large scale dynamicity in regulatory development, it has also been noted that divergence can appear at the TF binding level without altering overall regulatory signal (Venkataram and Fay, 2010). As a result, subtle differences in transcription factor binding may drive small but biologically meaningful differences in transcription factor binding affinity and combinatorial TF interactions between tissue types particularly utilising the broader repertoire of enhancer transcription factors.

Dynamicity in both chromatin accessibility and histone mark enrichment was statistically significant between developmental timepoints, suggesting that *cis-*regulatory turnover was a key driver of regulatory divergence across development. Chromatin accessibility showed the highest modulation, with promoter-associated ATAC displaying greater significant differential accessibility than enhancers across both species. However, this difference diminished at absolute log2 fold-change ≥ 1, indicating fewer but often greater changes in enhancer accessibility. Widespread promoter associated accessibility may not entirely correspond to similarly extensive differential gene expression as promoter accessibility alone is insufficient in determining transcriptional activity (Chereji *et al*., 2019). High difference in chromatin accessibility seems to coincide with positive correlation of differential histone mark enrichment in both promoters and enhancers, and has been previously shown that histone acetylation and methylation may lead to inhibition of higher chromatin structure formation leading to higher accessibility and transcription activity (Chen *et al., 2*024). However, *optix* and *cortex* promoter associated peaks showed a limited correlation between chromatin accessibility and histone mark enrichment, highlighting that accessibility is not strictly linked to changes in histone signal enrichment between developmental timepoints. Consequently, lower differential enrichment of H3K27ac enhancer compared to promoters may not be entirely representative of more stable enhancer activity. It has additionally been shown that diminishing H3K27ac marks at enhancers during mice embryonic development had limited impact on normal developmental mechanisms, likely due to chromatin accessibility and enrichment of other histone marks remaining unperturbed (Zhang *et al*., 2020). Together these patterns suggest that chromatin accessibility provides a more dynamic measure of developmental CRE state in this system, whereas histone modifications, particularly enhancer-associated H3K27ac, remain comparatively stable across temporal conditions.

### iv. Predicted enhancer regions enriched for CNEEs

Previously, it has been suggested that CNEEs likely contain conserved regulatory elements under evolutionary constraint, with putative regulatory function supported by enrichment of accessible chromatin within these regions (Edwards *et al*., 2017; Cicconardi *et al*., 2023). In Heliconiini, CNEEs overlapped ATAC-seq peaks at 2.4-fold and 2.0-fold enrichment above random expectation in brain and wing imaginal disc tissues respectively (Cicconardi *et al*., 2023). Here, we have highlighted significant enrichment of predicted *cis*-regulatory enhancer regions within CNEEs across 34 Heliconiini species, providing further support for this association and suggesting a substantial portion of conserved non-exonic sequences correspond to *cis*-regulatory regions. Note, however, that complete overlap is not expected, as CNEEs additionally encompass other functional elements, such as promoters, various non-coding RNAs, and other regulatory regions, and microproteins (Mohsen *et al*., 2023). Consistency of this enrichment is also suggestive of a conserved regulatory component maintained throughout the Heliconiini radiation. This is in line with previous work in Lepidoptera, demonstrating that CREs active across multiple developmental timepoints are markedly more conserved than stage specific elements (Lewis *et al*., 2016).

Despite this enrichment, correspondence between CNEEs and predicted enhancers was incomplete, with approximately 45.3% of CNEEs overlapping predicted enhancer regions and ∼32.3% of predicted enhancer regions overlapping CNEEs. Substantial proportions of enhancers lacking CNEE overlap may therefore present regulatory elements subject to weaker evolutionary constraint or lineage specific enhancer gains. These regions require further corroboration, as an absence of CNEE overlap does not necessarily indicate rapid regulatory evolution, but may instead reflect functional regulatory sequences lacking detectable conservation. However, this interpretation is consistent with comparative analyses in *D. melanogaster*, which display frequent enhancer gains between closely related species (Arnold *et al*., 2014). Further support for regulatory diversification in *Heliconius* has been validated via association of structural variations and differential chromatin accessibility, including transposable element insertions, suggesting a potential mechanism of lineage specific regulatory variation with additional recent studies demonstrating widespread regulatory divergence within the *H. erato* radiation (Ruggieri *et al*., 2024; Ruggieri *et al*., 2026). Additionally, conservation of CNEEs is itself not necessarily static with approximately 51% of Heliconiini CNEEs showing evidence of accelerated evolutionary rates at some point across the phylogeny (Cicconardi *et al*., 2023).

### v. Conclusion

This study provides a comprehensive annotation and characterisation of Heliconiini *cis*-regulatory dynamics and demonstrates the resultant datasets provide trends consistent with expected patterns of genomic distribution, sequence content, and CRE divergence. Promoters exhibited more conserved transcription factor binding architectures, whilst enhancers displayed greater flexibility and transcription factor motif repertoires. Additionally, *cis-*regulatory activity was highly conserved between hindwing and forewing tissues, suggesting that differences in wing patterning are unlikely to be primarily associated with differential CRE accessibility and histone enrichment alone. Instead, *cis-*regulatory divergence appears to be primarily temporal with substantial chromatin remodelling and differential histone enrichment highlighting CREs as a key mechanism underpinning transcriptional reprogramming across development. Extending these experimentally defined regulatory landscapes, sequence-based enhancer prediction demonstrated substantial cross species transferability facilitating enhancer annotation across 34 Heliconiini genomes. Predicted enhancers showed significant enrichment within independently characterised accessible chromatin and CNEE regions, identifying a potentially conserved component of the Heliconiini enhancer landscape alongside a substantial proportion of putative enhancers outside CNEEs that may provide greater regulatory flexibility. Collectively, these datasets provide a foundation for future comparative and functional investigations into enhancer conservation, regulatory turnover and their contributions to phenotypic diversification across the Heliconiini radiation.

## Data accessibility

Previously published sequencing datasets analysed in this study are publicly available through the NCBI Sequence Read Archive (SRA), with corresponding accession identifiers provided in Supplementary SeqData.tsv files. Code, supporting scripts, supplementary tables and quality control metrics generated during this study are publicly available through GitHub via: adamritchie4/heliconius-cis-regulatory-project

## Acknowledgments

This work was carried out using the computational facilities of the Advanced Computing Research Centre, University of Bristol. We are grateful to funding from the HFSP (RGP0029/2022) and Wellcome Trust Discovery Grant (312615/Z/24/Z).

