## Supplemental Materials: for "Characterisation of *cis*-regulatory elements and genome wide enhancer prediction in Heliconiini butterflies"

### Supplementary Materials

| Species | Lib | Tissue | Condition | SRRid | Replicate |
|---|---|---|---|---|---|
| Hlat | lib1 | HW | ATAC | SRR6653411 | rep1 |
| Hlat | lib2 | HW | ac | SRR6181407 | rep1 |
| Hlat | lib3 | HW | met | SRR6181408 | rep1 |
| Hlat | lib4 | HW | ATAC | SRR6653423 | rep2 |
| Hlat | lib5 | HW | ac | SRR6181431 | rep2 |
| Hlat | lib6 | HW | met | SRR6181432 | rep2 |
| Hlat | lib7 | FW | ATAC | SRR6653410 | rep1 |
| Hlat | lib8 | FW | ac | SRR6181405 | rep1 |
| Hlat | lib9 | FW | met | SRR6181406 | rep1 |
| Hlat | lib10 | FW | ATAC | SRR6653422 | rep2 |
| Hlat | lib11 | FW | ac | SRR6181429 | rep2 |
| Hlat | lib12 | FW | met | SRR6181430 | rep2 |
| Hhim | lib25 | HW | ATAC | SRR6653407 | rep1 |
| Hhim | lib26 | HW | ac | SRR6181399 | rep1 |
| Hhim | lib27 | HW | met | SRR6181400 | rep1 |
| Hhim | lib28 | HW | ATAC | SRR6653419 | rep2 |
| Hhim | lib29 | HW | ac | SRR6181423 | rep2 |
| Hhim | lib30 | HW | met | SRR6181424 | rep2 |
| Hhim | lib31 | FW | ATAC | SRR6653406 | rep1 |
| Hhim | lib32 | FW | ac | SRR6181397 | rep1 |
| Hhim | lib33 | FW | met | SRR6181398 | rep1 |
| Hhim | lib34 | FW | ATAC | SRR6653418 | rep2 |
| Hhim | lib35 | FW | ac | SRR6181421 | rep2 |
| Hhim | lib36 | FW | met | SRR6181422 | rep2 |
| Hhim | lib37 | FW | input | SRR6181445 | control |
| Hhim | lib38 | HW | input | SRR6181446 | control |
| Hlat | lib39 | FW | input | SRR6181449 | control |
| Hlat | lib40 | HW | input | SRR6181450 | control |

**Table S1.** D3SeqData.tsv sequencing dataset information for day 3 post-pupation (D3) *Heliconius himera* and *H. erato lativitta* forewing (FW) and hindwing (HW) tissues. Datasets comprise ATAC-seq, H3K27ac (ac) and H3K4me3 (met) ChIP-seq, with two biological replicates per assay and tissue, alongside corresponding ChIP-seq input controls. Species, library identifiers, tissue, assay conditions, Sequence Read Archive (SRA) accession identifiers and replicate information are provided for each dataset.

| Species | Lib | Tissue | Condition | SRRid |
|---|---|---|---|---|
| Hhim | lib32 | FW | ac | SRR6181401 |
| Hhim | lib33 | FW | met | SRR6181402 |
| Hhim | lib26 | HW | ac | SRR6181403 |
| Hhim | lib27 | HW | met | SRR6181404 |
| Hlat | lib8 | FW | ac | SRR6181409 |
| Hlat | lib9 | FW | met | SRR6181410 |
| Hlat | lib2 | HW | ac | SRR6181411 |
| Hlat | lib3 | HW | met | SRR6181412 |
| Hhim | lib35 | FW | ac | SRR6181425 |
| Hhim | lib36 | FW | met | SRR6181426 |
| Hhim | lib29 | HW | ac | SRR6181427 |
| Hhim | lib30 | HW | met | SRR6181428 |
| Hlat | lib11 | FW | ac | SRR6181433 |
| Hlat | lib12 | FW | met | SRR6181434 |
| Hlat | lib5 | HW | ac | SRR6181435 |
| Hlat | lib6 | HW | met | SRR6181436 |
| Hhim | lib31 | FW | ATAC | SRR6653408 |
| Hhim | lib25 | HW | ATAC | SRR6653409 |
| Hlat | lib7 | FW | ATAC | SRR6653412 |
| Hlat | lib1 | HW | ATAC | SRR6653413 |
| Hhim | lib34 | FW | ATAC | SRR6653420 |
| Hhim | lib28 | HW | ATAC | SRR6653421 |
| Hlat | lib10 | FW | ATAC | SRR6653424 |
| Hlat | lib4 | HW | ATAC | SRR6653425 |

**Table S2.** OMSeqData.tsv sequencing dataset information for day 7 ommochrome (OM) stage *Heliconius himera* and *H. erato lativitta* forewing (FW) and hindwing (HW) tissues. Datasets comprise ATAC-seq, H3K27ac (ac) and H3K4me3 (met) ChIP-seq, with two biological replicates per assay and tissue, alongside corresponding ChIP-seq input controls. Species, library identifiers, tissue, assay conditions, Sequence Read Archive (SRA) accession identifiers and replicate information are provided for each dataset.

| Species | Lib | Tissue | Sample | SRRid |
|---|---|---|---|---|
| Hcha | lib1 | Head | CH_2 | SRR17466641 |
| Hcha | lib2 | Head | CH_6 | SRR17466640 |
| Hcha | lib3 | FW | CH_FW_2 | SRR17466627 |
| Hcha | lib4 | FW | CH_FW_7 | SRR17466626 |
| Hcha | lib5 | HW | CH_HW_2 | SRR17466637 |
| Hcha | lib6 | HW | CH_HW_7 | SRR17466636 |
| Hdem | lib7 | Head | BR_12 | SRR17466631 |
| Hdem | lib8 | Head | BR_14 | SRR17466630 |
| Hdem | lib9 | FW | LI_FW_7 | SRR17466625 |
| Hdem | lib10 | FW | pboy | SRR17466624 |
| Hdem | lib11 | HW | LI_HW_7 | SRR17466635 |
| Hdem | lib12 | HW | LB_42 | SRR17466634 |
| Hmel | lib13 | Head | BR_11 | SRR17466629 |
| Hmel | lib14 | Head | M4 | SRR17466628 |
| Hmel | lib15 | FW | LI_FW_14 | SRR17466639 |
| Hmel | lib16 | FW | M4_FW | SRR17466638 |
| Hmel | lib17 | HW | LI_HW_14 | SRR17466633 |
| Hmel | lib18 | HW | M4_HW | SRR17466632 |

**Table S3.** ATAC-seq datasets used for independent validation of CNN-predicted enhancers across *Heliconius* lineages. Previously published ATAC-seq datasets including *H. charithonia*, *H. e. demophoon* and *H. melpomene*, representing the sara-sapho, erato and melpomene lineages respectively, were used to independently validate predicted enhancer regions across phylogenetically divergent taxa. Datasets comprised head, forewing (FW) and hindwing (HW) tissues, with two biological samples per tissue and species. Species, library identifiers, tissue, sample identifiers and Sequence Read Archive (SRA) accession numbers are provided for each dataset.

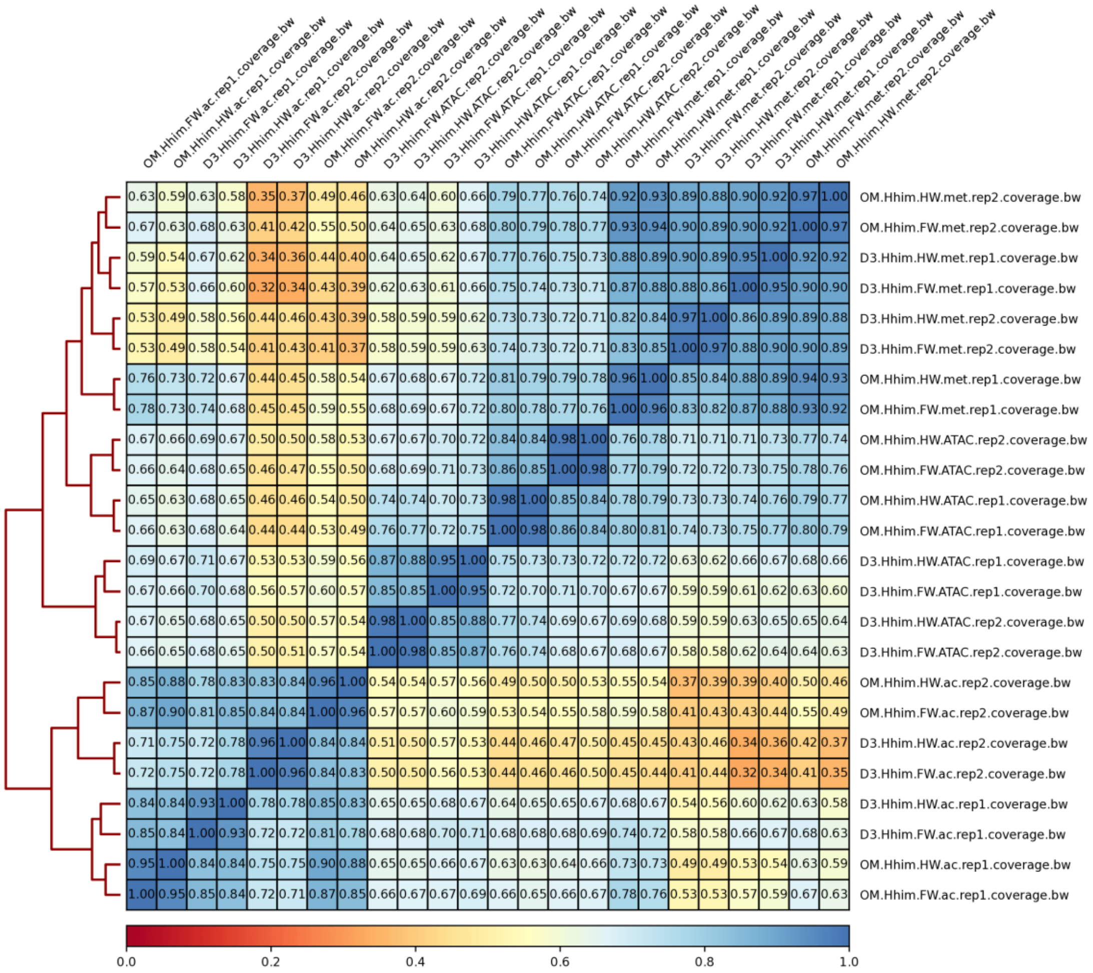


**Figure S1.** Replicate concordance across conditions in *Heliconius himera*. Heatmap displays pairwise correlations between biological replicates across ATAC-seq, H3K27ac, and H3K4me3 ChIP-seq datasets.

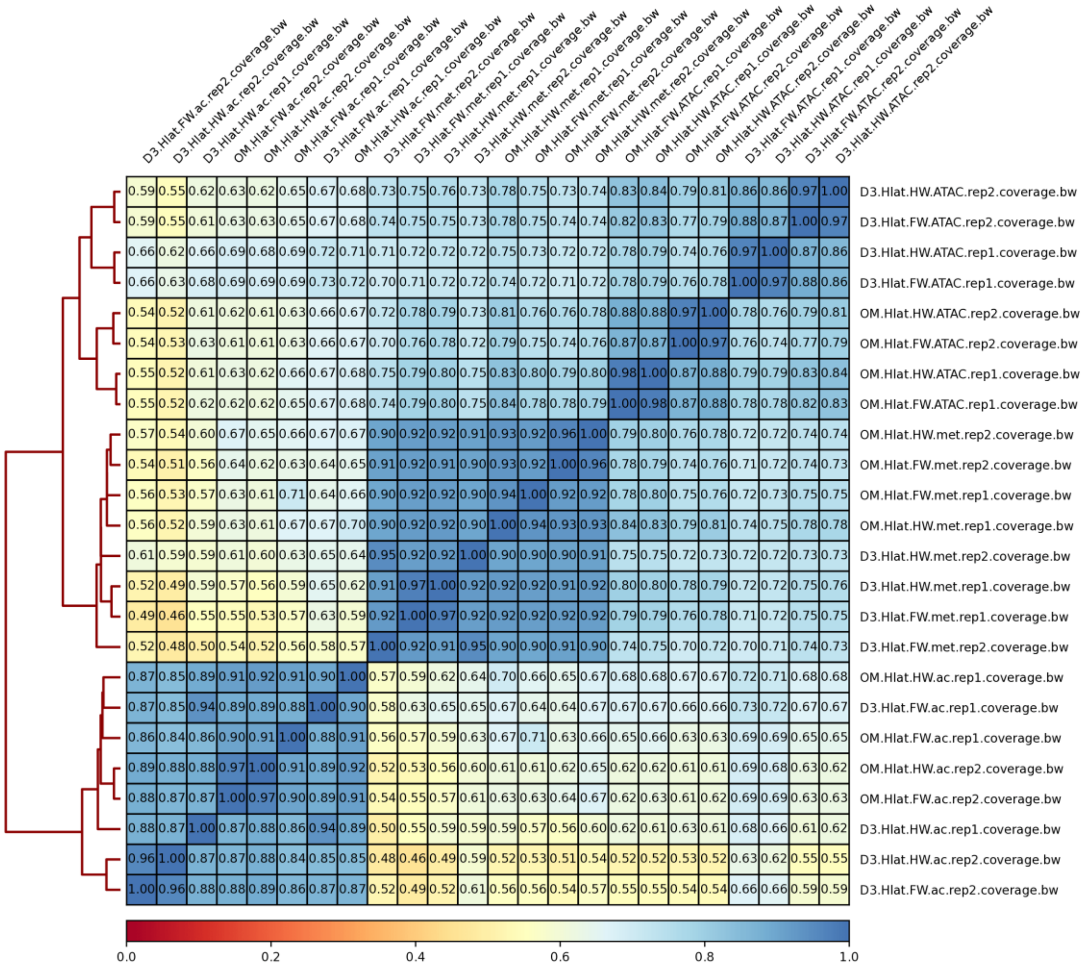


**Figure S2.** Replicate concordance across conditions in *Heliconius erato lativitta*. Heatmap displays pairwise correlations between biological replicates across ATAC-seq, H3K27ac, and H3K4me3 ChIP-seq datasets.

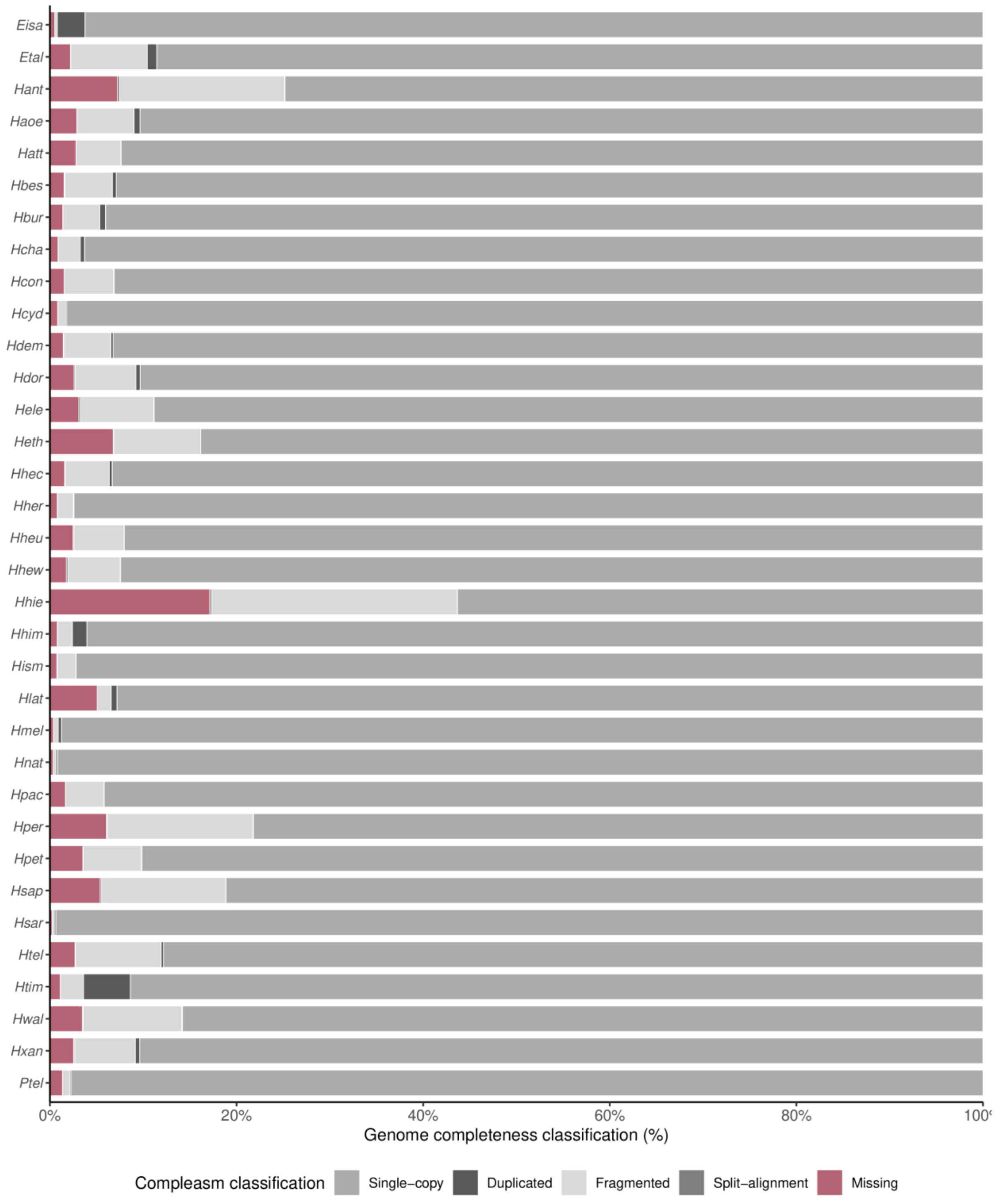


**Figure S3.** Genome assembly completeness across 34 Heliconiini species. Stacked bar chart displaying genome assembly completeness scores assessed using Compleasm v0.2.10 with the lepidoptera_odb12 lineage dataset. Orthologs were classified as single-copy (S), duplicated (D), fragmented (F), split-alignment (I) or missing (M). Bars represent the percentage of orthologs assigned to each classification for their respective species.

#### A note on *Heliconius erato petiverana*

An initial HW-D3 condition-specific pipeline and analytical framework applied to *H. himera* and *H. e. lativitta* was additionally applied to *Heliconius erato petiverana* for annotation of *cis*-regulatory promoters and enhancers. Genome quality assessment indicated completeness of the *H. e. petiverana* assembly via compleasm at 90.1%, whilst contiguity analysis identified an assembly structure with an N50 of 8.53 Mb and N90 of 1.90 Mb across a 318.0 Mb assembly. Whilst the assembly exhibited reduced completeness compared to *H. himera* and *H. e. lativitta*, these metrics still warranted inclusion of *H. e. petiverana* in downstream analysis.

Following identical preprocessing, quality assessment, and peak calling procedures, preliminary condition-specific analysis of *H. e. petiverana* generated a comparable 92,352 ATAC peaks, 44,988 H3K27ac peaks, and finally 30,127 H3K4me3 peaks in D3-HW samples. Initial classification relied exclusively on ATAC and histone peak overlap, with promoters defined by ATAC, H3K27ac and H3K4me3 enrichment, and enhancers by ATAC and H3K27ac enrichment in the absence of H3K4me3, without additional TSS-based classification requirements. Intersection of these datasets identified 10,465 merged putative promoters, and 32,938 merged putative enhancers, displaying an increased capture of *cis*-regulatory elements in *H. e. petiverana* compared to *H. himera* and *H. e. lativitta*. However, subsequent characterisation and regulatory mapping of these *cis*-regulatory elements and peak characteristics revealed several deviations from the patterns observed in *H. himera* and *H. e. lativitta*. Initial quality assessment identified aberrant ATAC signal profiles in *H. e. petiverana*, with 1 ATAC-seq sample exhibiting a right-shifted fragment size distribution indicating increased mononucleosome fragments compared to the expected enrichment of nucleosome-free regions. Similarly, one H3K4me3 replicate (D3 forewing replicate 2) displayed reduced signal quality (FRiP = 0.06). Across the merged datasets, *H. e. petiverana cis*-regulatory elements additionally displayed increased ATAC-seq signal intensity and reduced H3K4me3 signal intensity relative to *H. himera* and *H. e. lativitta*, consistent with the patterns observed during data preprocessing. Finally, H3K4me3 peak size distributions displayed altered characteristics, with H3K4me3 peaks showing reduced representation of broader peak regions (H3K4me3 median peak size = 238 bp) (Figure S4).

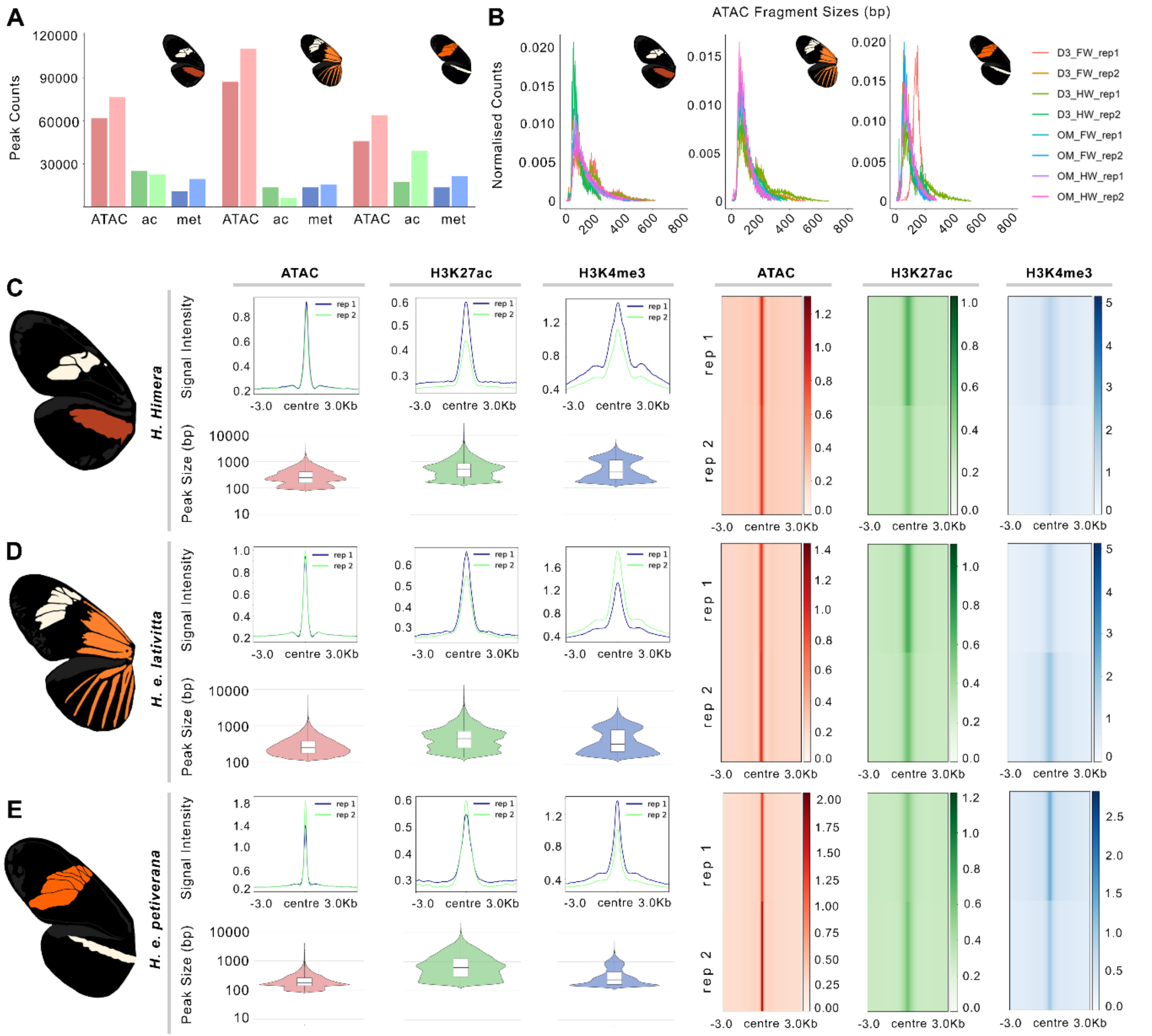


**Figure S4.** Preprocessing and peak calling quality checks for *Heliconius* data. (A) Overview of total peak counts for ATAC, H3K27ac, and H3K4me3 marks per species. (B) ATAC fragment size analysis for each species confirming high quality ATAC seq profiles. (C-E) Genome wide peak calling quality checks for each *Heliconius* species, showing coverage profiles in peak centred 3kb windows and heatmaps for each sequencing mark, alongside peak size distribution boxplots of all peaks used in downstream analysis.

The apparent technical constraints appeared to increase ambiguity in putative promoter and enhancer classification. From early condition-specific D3-HW testing, whilst promoter-enhancer overlap between *cis*-regulatory elements using no TSS-based classification requirement remained low in *H. himera* (3.9%) and *H. e. lativitta* (2.3%), *H. e. petiverana* displayed a substantial increase in promoter-enhancer dataset overlap at 17.3%. This overlap appeared to cause downstream complications, with enhancer-associated regions showing increased promoter-proximal enrichment, however this trend was additionally noted in the replicate supported datasets which displayed no overlapping *cis*-regulatory elements. Putative promoter enrichment was additionally reduced in *H. e. petiverana* at promoter regions in comparison to *H. himera* and *H. e. lativitta* (Figure S5). These results suggested that the aberrant ATAC profiles and reduced H3K4me3 peak sizes may have contributed to increased ambiguity between the enhancer and promoter classifications.

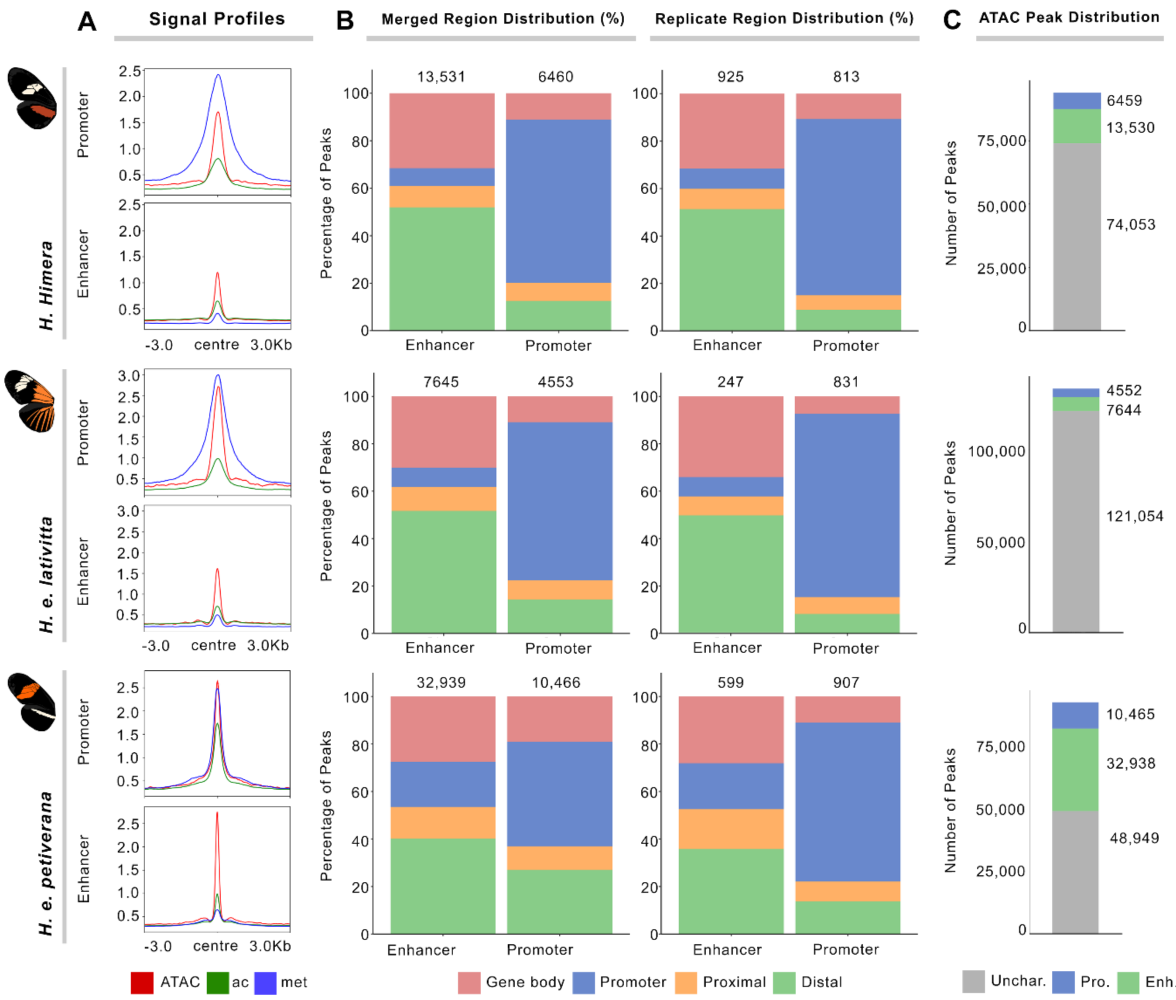


**Figure S5.** Promoter and enhancer dataset creation summary. (A) Displays signal profiles for ATAC, H3K27ac (ac), and H3K4me3 (met) present in each species promoter and enhancer BED files with signal intensity on y-axis and peaks 3kb around centres on x-axis. (B) Displays *cis*-regulatory mapping plots for promoters and enhancers per species (%), merged datasets represents union on all CREs in each replicate, whilst replicate datasets represent union of only CREs present in both replicates. (C) Displays overall distribution of CREs across accessible chromatin landscape per species (%).

Following this preliminary testing, the classification framework was refined to additionally require promoters to occur within 1 kb of annotated transcription start sites (TSSs) and enhancers outside these regions, alongside the existing ATAC and histone enrichment requirements. This refined classification was subsequently adopted for the main comparative analyses of *H. himera* and *H. e. lativitta*. Whilst the additional TSS-based restrictions reduced the apparent ambiguity between regulatory classifications, they also obscured some of the underlying inconsistencies identified during preliminary testing. Subsequent analysis of *H. e. petiverana* using the refined framework continued to display deviations in downstream regulatory characteristics, suggesting that the underlying data quality constraints remained unresolved.

These deviations were also apparent at key wing patterning loci *optix* and *cortex*. At the *cortex* locus, a promoter-associated region was identified within the expected transcription start site region, however no CRE association was observed for *optix*. Conversely, distal enhancer associations surrounding *optix* and *cortex* were elevated compared with *H. himera* and *H. e. lativitta*, corresponding with the increased number of identified CREs in *H. e. petiverana*. While this may reflect genuine regulatory complexity, the magnitude of increased CRE identification suggests that technical factors likely contributed to elevated regulatory element capture (Figure S6).

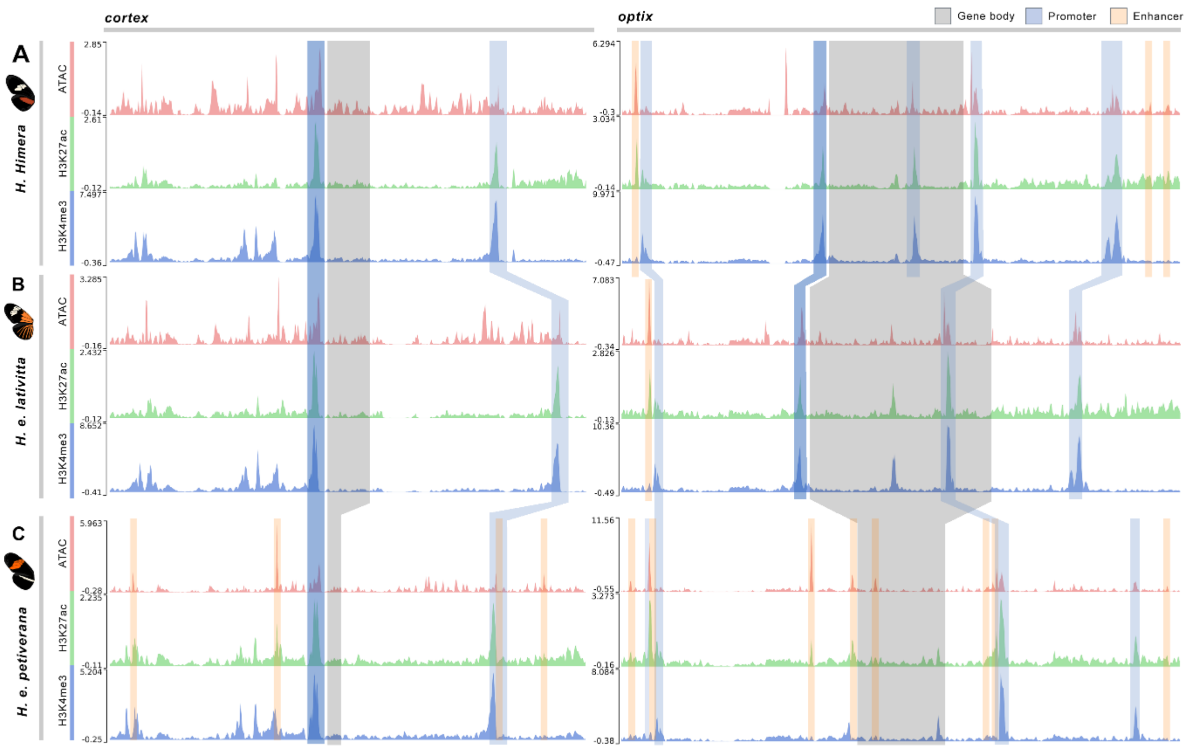


**Figure S6.** Signal track profiles in 30kb windows around the *cortex* and *optix* target genes including relative intensity scores for each signal track per gene and species. (A-B) Indicate successful capture of cortex and optix cognate promoters alongside additional CREs within the 30kb window. (C) Presents successful capture of cortex promoter, though optix promoter appears to be misclassified as an enhancer due to truncation of the H3K4me3 peak.

Further deviation was observed in motif and differential accessibility analyses. Whilst GC content distributions remained comparable between the three species, motif clustering patterns deviated from the patterns in *H. himera* and *H. e. lativitta* which both coincided with expectations from the literature. Promoter and enhancer motif clustering displayed a consistent and clear divide within *H. himera* and *H. e. lativitta*, with enhancers displaying greater motif diversity than promoters, however clustering ratios between *H. e. petiverana* promoters and enhancers exhibited similar profiles. This pattern is consistent with the increased ambiguity and overlap in promoter and enhancer classification. Similarly, differential transcription factor binding analyses displayed reduced promoter-specific motif enrichment and increased enhancer-associated signal shifts compared with the other species (Figure S7).

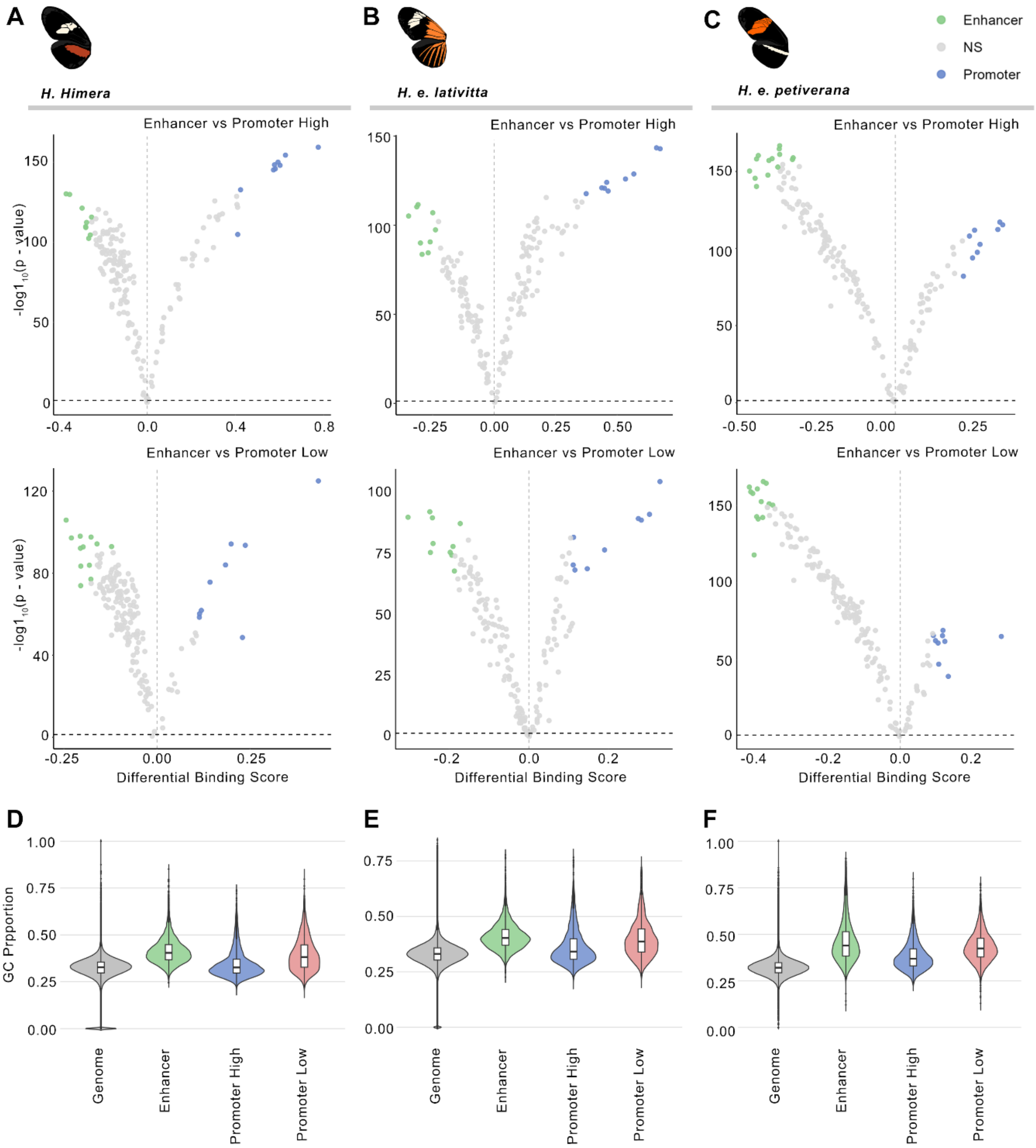


**Figure S7.** Displays summary of differential motif binding analysis between CRE datasets alongside GC enrichment. (A-C) Displays differential log fold change binding scores on x-axis of motifs identified in promoter and enhancer datasets per species with negative log of p-value on y-axis, indicating clear shift towards conserved high affinity binding motifs in high confidence promoters and more broad distribution of enhancer motifs in *H. himera* and *H. e. lativitta*. (D-F) Display GC content analysis for genome wide background, enhancers, high confidence promoters, and low confidence promoters per species (%).

Finally, in contrast to the high degree of histone enrichment and chromatin accessibility profile similarities observed in *H. himera* and *H. e. lativitta*, *H. e. petiverana* continued to display aberrant profile characteristics. H3K27ac enrichment dynamicity was observed in enhancer tissue comparisons with significant downregulation of hindwing marks compared to forewing. Additionally, dual clustering of *H. e. petiverana* H3K4me3 signal in promoters and dropout of tissue comparison H3K27ac peaks alongside previously noted inconsistencies in data preprocessing suggest these are unlikely to represent true biological trends and may instead reflect data quality constraints (Figure S8).

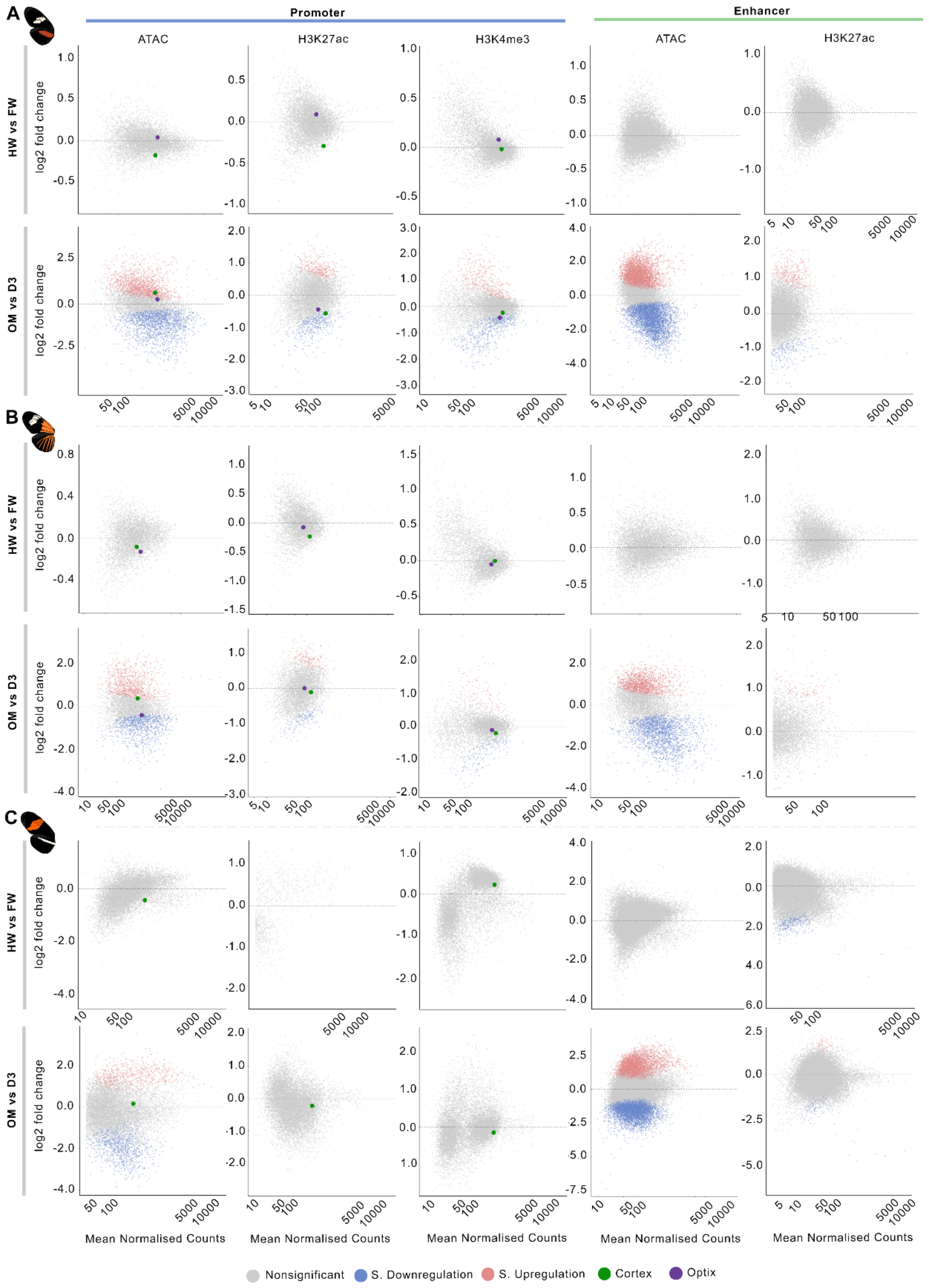


**Figure S8.** Displays differential enrichment analysis of signal types corresponding to promoters and enhancer across spatiotemporal conditions with $\log_2$ fold change displayed on y-axis and mean normalised counts per signal type on y-axis. (A-B) H. himera and H. e. lativitta display highly conserved trends of minimal differential expression between hindwing and forewing tissue types, with considerably higher dynamicity between temporal development. (C) Differential enrichment associated with H. e. petiverana marks exhibiting aberrant profiles. Promoter marks associated with cortex and optix genes additionally plotted for visual reference

Due to the observed data constraints in *H. e. petiverana* chromatin profiles, histone peak morphology, and regulatory class delineation, biological interpretation of *H. e. petiverana cis*-regulatory landscapes was assigned reduced confidence. Whilst the refined TSS-based classification reduced the visibility of some of the initial classification ambiguities, persistence of downstream inconsistency suggested that underlying technical artifacts remained unresolved. Consequently, comparative analysis was focused on the promoter and enhancer characteristics of *H. himera* and *H. e. lativitta*, both for biological interpretation and machine learning conservation training data, whilst *H. e. petiverana* results are presented as supplementary evidence highlighting both the applicability and limitations of comparative *cis*-regulatory element annotation via an ATAC and histone mark system.
